# Retinoic acid control of pax8 during renal specification of Xenopus pronephros involves hox and meis3

**DOI:** 10.1101/2022.06.21.496994

**Authors:** Jennifer Durant-Vesga, Nanoka Suzuki, Haruki Ochi, Ronan Le Bouffant, Alexis Eschstruth, Hajime Ogino, Muriel Umbhauer, Jean-François Riou

## Abstract

Development of the Xenopus pronephros relies on renal precursors grouped at neurula stage into a specific region of dorso-lateral mesoderm called the kidney field. Formation of the kidney field at early neurula stage is dependent on retinoic (RA) signaling acting upstream of renal master transcriptional regulators such as pax8 or lhx1. Although *lhx1* might be a direct target of RA-mediated transcriptional activation in the kidney field, how RA controls the emergence of the kidney field remains poorly understood. In order to better understand RA control of renal specification of the kidney field, we have performed a transcriptomic profiling of genes affected by RA disruption in lateral mesoderm explants isolated prior to the emergence of the kidney field and cultured at different time points until early neurula stage. Besides genes directly involved in pronephric development (*pax8, lhx1, osr2, mecom*), hox (*hoxa1, a3, b3, b4, c5* and *d1*) and the hox co-factor *meis3* appear as a prominent group of genes encoding transcription factors (TFs) downstream of RA. Supporting the idea of a role of meis3 in the kidney field, we have observed that meis3 depletion results in a severe inhibition of *pax8* expression in the kidney field. Meis3 depletion only marginally affects expression of *lhx1* and *aldh1a2* suggesting that meis3 principally acts upstream of *pax8*. Further arguing for a role of meis3 and hox in the control of pax8, expression of a combination of meis3, hoxb4 and pbx1 in animal caps induces *pax8* expression, but not that of *lhx1*. The same combination of TFs is also able to transactivate a previously identified *pax8* enhancer, Pax8-CNS1. Mutagenesis of potential PBX-Hox binding motifs present in Pax8-CNS1 further allows to identify two of them that are necessary for transactivation. Finally, we have tested deletions of regulatory sequences in reporter assays with a previously characterized transgene encompassing 36.5 kb of the *X. tropicalis pax8* gene that allows expression of a truncated pax8-GFP fusion protein recapitulating endogenous *pax8* expression. This transgene includes three conserved *pax8* enhancers, Pax8-CNS1, Pax8-CNS2 and Pax8-CNS3. Deletion of Pax8-CNS1 alone does not affect reporter expression, but deletion of a 3.5kb region encompassing Pax8-CNS1 and Pax8-CNS2 results in a severe inhibition of reporter expression both in the otic placode and kidney field domains.

## Introduction

The nephron is the basic unit of vertebrate kidneys. Although renal function can be very different in animals living in fresh water and in terrestrial animals, the general organization of the nephron is remarkably conserved (Desgrange and Cereghini, 2015). In fishes and amphibia, the pronephros is the functional kidney at larval stages. The Xenopus tadpole bears a pair of pronephroi located on each side of the body that are made of one single giant nephron (Wessely and Tran, 2011). This nephron is filtering plasma at the level of the glomus, while filtrate collected in the nephrocoele is further transformed into definitive urine by reabsorption of ions and proteins in the tubule. Notably, the different cell types constituting the Xenopus pronephros are very similar to those of the metanephric nephron of mammals, as evidenced by the conserved expression of orthologs of genes of the solute-carrier and claudin gene families for tubular cell types (Raciti et al., 2008), and podocin or nephrin expressed in podocytes of the glomus (White et al., 2010). Understanding the molecular mechanisms controlling specification of renal precursors of the Xenopus pronephros is therefore not only of interest for the study of vertebrate kidney development. It can also provide valuable information in the perspective of renal tissue engineering (Lienkamp, 2016).

Renal precursors at the origin of the pronephros in Xenopus are grouped in a specific region of dorso-lateral mesoderm arising at the onset of neurulation. This region called the kidney field is defined by the overlapping expression of *pax8* and *lhx1* genes (Carroll and Vize, 1999). Renal precursors of the kidney field also express *osr1* and *osr2* genes (Tena et al., 2007). These four different genes encode transcription factors (TF), all of which have been shown to be required for pronephric development (Buisson et al., 2015; Cirio et al., 2011; Tena et al., 2007). We have previously shown that pax8 plays a critical role for the specification of tubular cell types. Pax8 depletion causes an inhibition of the formation of the pronephric tubule anlage at early tailbud stage, resulting in the total absence of the tubule in the tadpole. At neurula stages, expression of *lhx1* and *osr2* in the kidney field is not affected, but *hnf1b* expression is inhibited. Pax8 depletion is also causing cell proliferation defects in the kidney field in a canonical-wnt-dependent manner, potentially through the control of components of the wnt signaling pathway such as *dvl1* or *sfrp3* (Buisson et al., 2015).

Mechanisms controlling the emergence of the kidney field are only partially understood. Wnt11b can induce pronephric structures in unspecified lateral mesoderm explants, and could act as a potential inducer (Tetelin and Jones, 2010). However, retinoic acid (RA) signaling appears to be absolutely required. Its disruption causes a loss of *pax8* and *lhx1* expression in the kidney field, resulting in the inhibition of pronephric markers of tubule and glomus at tailbud stages, and eventually the absence of pronephros in the tadpole (Cartry et al., 2006). Control of *pax8* and *lhx1* expression in the kidney field by RA is still poorly understood. Some clues indicate that control of *pax8* and *lhx1* involves different mechanisms. For example, up-regulation of *lhx1* in response to exogenous RA is very fast both in embryos (Cartry et al., 2006) and animal caps treated with activin and RA (Drews et al., 2011), while *pax8* response is delayed. *Lhx1* response to RA treatment also occurs in the absence of protein synthesis, supporting the idea that *lhx1* might be a direct target of RA-mediated transcriptional activation in the kidney field (Cartry et al., 2006; Drews et al., 2011). *Pax8* expression in the kidney field involves TRPP2-dependent calcium signaling, which requires RA potentially through the control of TRPP2 trafficking to the plasma membrane (Futel et al., 2015). However, this mechanism is likely to act in a permissive rather than in an instructive way, suggesting that more RA-dependent inputs control the expression of *pax8* in the kidney field. Regulation of *pax8* expression in the *Xenopus* embryo is taking place through a set of evolutionary conserved enhancers associated with a silencer located in the *pax8* proximal promoter sequence. This silencer suppresses enhancer activity outside *pax8* expression domain (Ochi et al., 2012). How these regulatory elements are potentially used to control *pax8* expression in the kidney field, and what are the inputs is largely elusive.

In order to better understand the transcriptional response to RA during pronephric precursor specification, we have performed expression profiling of RA target genes expressed in lateral mesoderm at the origin of the kidney field. One prominent group of genes involved in the transcriptional response to RA corresponded to hox family members and the hox cofactor meis3. Meis3 depletion in ventro-lateral mesoderm results in a severe inhibition of *pax8* expression in the kidney field, suggesting it is part of the mechanisms acting upstream of *pax8*. We further provide results suggesting that meis3 is involved in the regulation of *pax8* in combination with hox. First, we show that a combination of hoxb4 with pbx1 and meis3 cofactors is able to upregulate *pax8* in isolated animal caps. We further show that the same combination is able to transactivate one conserved enhancer of the *pax8* gene, Pax8-CNS1 (Ochi et al., 2012). Finally, we have identified a 3.5kb region of the *pax8* gene that is required for *pax8* expression at neurula stage both in the kidney field and the otic placod. This region contains two conserved *pax8* enhancers, Pax8-CNS1 and Pax8-CNS2. These results point to hox/meis3 are part of the complex regulation of *pax8* during renal specification of the pronephros.

## Materials and methods

### Embryos, microinjection and microdissections

*Xenopus* embryos have been obtained as previously described by human chorionic gonadotropin stimulation of females and in vitro fertilization (Futel et al., 2015). They were cultured in modified Barth’s Solution (MBS) and staged according to (Nieuwkoop and Faber, 1975). All animal experiments were carried out according to approved guidelines validated by the Ethics Committee on Animal Experiments “Charles Darwin”(C2EA-05) with the “Autorisation de projet” number 02164.02.

Microinjection of morpholinos and mRNA was performed as described in 0.1x MBS containing 3% Ficoll (Colas et al., 2008). Synthesis of capped mRNA was done as previously described (Umbhauer et al., 2000). Dissections of blastula animal caps and gastrula LMZ explants were carried out in 1xMBS in agar-coated dishes using platinum wires and loops. Culture of explants was carried out in 1xMBS. For LMZ dissection, care was taken to exclude blastocoel wall in order to avoid contamination by ectodermal tissue, as previously described (Le Bouffant et al., 2012).

### Constructs

*X. laevis hoxb4.S* and *pbx1.L* coding sequences were cloned in pCRII vector (InVitrogen) by RT-PCR amplification from NFst28 cDNA. *X. laevis meis3.S* coding sequence was PCR-amplified from a clone kindly provided by Dr AH Monsoro. *Hoxb4.S, pbx1.L* and *meis3.S* coding sequences were inserted into pCS2+ vectors for RNA synthesis or expression in HEK293 cells. PCR mutagenesis has been performed to introduce silent mutations downstream of the start codon in order to avoid binding to Momeis3 in rescue experiments (AUG GCA CAA AGG to AUG GC**T** CA**G CGC**). Use of p-βSRN3-GFP (ZernickaGoetz et AL., 1996) and pCS2XCyp26 (Hollemann et al., 1998) plasmids has been previously described (Le Bouffant et al., 2012). For transactivation assays, *X. tropicalis* Pax8-CNS1 sequence was removed from the 1^pax8^-pax8(-2038)-GFP construct (Ochi et al., 2012) with bamH1 and inserted upstream of the basal promoter of pGL4.23[luc2]miniP (PROMEGA) to generate pax8CNS1-luc. Mutations of Pbx-Hox and Meis/TGIF motifs in pax8CNS1-luc (pax8CNS1mut-lucMeis, pax8CNS1mut-lucHP1, pax8CNS1mut-lucHP2, pax8CNS1mut-lucHP3, pax8CNS1mut-lucHP12, pax8CNS1mut-lucHP13, pax8CNS1mut-lucHP23, pax8CNS1mut-lucHP123) were obtained with the QuikChange^®^ II XL Site-Directed Mutagenesis Kit (Agilent Technologies) using primers listed in Supplementary Table 4. Momeis3 antisense morpholino meis3 5’-CCTTTGTGCCATTCCGAGTTGGGTC-3’ has been previously characterized (In der Rieden et al., 2011).

### Microarray analysis

Embryos were microinjected equatorially at the 4-cell stage into the four blastomeres with 10 nl each of a mixture of cyp26a1 mRNA (25pg/nl) and GFP mRNA (50pg/nl), or with GFP mRNA alone (50pg/nl). LMZ explants were dissected immediately after appearance of the dorsal blastoporal lip and were kept at 14°C during one hour for healing. At this stage siblings were at NFst10-10.25. Explants were further cultured for 3h at 20°C (siblings at NFst11) and then placed overnight at 14°C until siblings reached early neurula stage (NFst14). Batches of 15 explants were frozen at each stage in 300 μl RLT solution (Qiagen) for further RNA extraction with Qiagen RNeasy columns. Experiments were validated for microarray analysis after RT-qPCR control of equal proportions of endoderm (*sox17β*) and mesoderm (*eomes*) in control and cyp26a1 gastrula stage samples, as well as minimal contamination of neural tissue (*sox2*). Proper RA disruption was controlled by scoring inhibition of pronephric markers *pax8* and *lhx1 in* neurula stage samples as previously described (Le Bouffant et al., 2012).

Expression profiling was performed on GeneChip™ Xenopus laevis Genome 2.0 Array (Affymetrix) at the Curie Institute Affymetrix platform. Probe Set analysis was conducted in R (R Core Team, 2019). Data from .CEL files (GEO accession number GSE205827) was read and normalized using the “affy” package (Gautier et al., 2004) to create the expression set. Differential expression analysis was further performed using “limma” package (Ritchie et al., 2015). Batch effect between the three analyzed experiments was adjusted by setting eBayes function’s argument robust to True (Phipson et al., 2016). Xenopus gene identification of Affymetrix probe Set IDs can be found at: http://ftp.xenbase.org/pub/GenePageReports/GenePageAffymetrix_laevis2.0_chd.txt

Gene annotation from Ensembl databases was retrieved using “biomaRt” R package (Durinck et al., 2009). In several instances where gene annotation from Ensembl was missing, annotation was added manually from Xenbase gene pages (http://www.xenbase.org/).

### Real time quantitative PCR and whole mounted in situ hybridization

Real time quantitative PCR (RT-qPCR) has been performed as previously reported (Le Bouffant et al., 2012). The Comparative Ct method was used to determine the relative quantities of mRNA, using *ornithine decarboxylase (odc*) or *β-actin* mRNA as endogenous reporter. Every RNA sample was analyzed in duplicate. Data point represents the mean ± SEM of at least three independent experiments. Data were analyzed using R by paired Student’s t-test. Primers used for RT-qPCR are given in supplementary table 4.

Whole mounted ISH has been carried out as previously described (Colas et al., 2008) with riboprobes for *pax8, lhx1* (Carroll and Vize, 1999), *aldh1a2* (Chen et al., 2001), *cyp26a1* (Hollemann et al., 1998) and *GFP* (Ochi et al., 2012).

### Transactivation experiments in HEK 293 cells

HEK293 cells were cultured in Dulbecco’s modified Eagle’s medium (DMEM) supplemented with 10% fetal calf serum. They were plated into 12-well plates at a density of 2.5 × 10^5^ cells/well. After 24 hours transfection, mixtures containing Opti-MEM^®^ Reduced Serum Media (ThermoFisher), 500ng of each reporter vector (pax8CNS1-luc, pax8CNS1mut-lucMeis, pax8CNS1mut-lucHP1, pax8CNS1mut-lucHP2 or pax8CNS1mut-lucHP3, pax8CNS1mut-lucHP12, pax8CNS1mut-lucHP13, pax8CNS1mut-lucHP23, pax8CNS1mut-lucHP123), 5 ng of pCMV-renilla luciferase normalization vector and the transactivation vector mix (pCS2-hoxb4 and/or pCS2-meis3 and/or pCS2-pbx1 and pCS2+ empty vector− 300ng total) were prepared. Subsequently, cells were transfected with these mixtures using 2.4μL of X-tremeGENE HP DNA Transfection Reagent (Roche) according to the manufacturer’s protocol. Transfected cells were washed twice in PBS, followed by the addition of 200μl 1X passive lysis buffer (Promega, Madison, WI, USA). All values are shown as the mean ± SEM. All determinations were repeated at least three times. Statistical significance was using Dunn’s multiple comparison test after Kruskal-Wallis test.

### Transgenic reporter assay

Transgenic *Xenopus* embryos carrying the wild-type *pax8*-exon2-GFP construct (previously described as *pax8*-GFP fosmid) were generated as previously described (Ochi et al., 2012) by sperm nuclear transplantation method with oocyte extracts (Hirsch et al., 2002; Kroll and Amaya, 1996). A mutant construct lacking CNS1 (Δ-CNS1) was generated from this wild-type construct by replacing the CNS1 with a *kan* gene cassette amplified from an EGFP-flpe-KAN plasmid using homologous recombination technique (DeLaurier et al., 2010), followed by flipping out of the *kan* cassette by arabinose-induced Flpe recombinase (Lee et al., 2001). A mutant construct lacking both CNS1 and CNS2 (Δ-CNS1/2) was generated by replacing a region encompassing CNS1 and CNS2 with the *kan* gene cassette as with the Δ-CNS1 construct. The *kan* cassette was not removed by the Flpe recombinase from this Δ-CNS1/2 construct. The manipulated embryos were cultured until the late neurula stage (NFst18/19), and all normally developed embryos were subjected to in situ hybridization to examine their GFP expression with maximum sensitivity (Ochi et al., 2012).

## Results

### Expression profiling of RA targets in lateral marginal zone explants

Active RA signaling is thought to be required for pronephric development during gastrulation and early neurulation, since treatment with a RAR inhibitor has only limited effects on pronephric development when it is applied after completion of gastrulation (Cartry et al., 2006). During gastrulation, cells that will give rise to renal precursors of the kidney field are localized in the lateral marginal zone (LMZ). In order to identify RA targets potentially involved in renal precursors specification, especially those involved in its transcriptional control, we have performed expression profiling of genes modified by RA disruption in LMZ explants isolated at early gastrula stage (NFst10) (Le Bouffant et al., 2012). LMZ explants taken from embryos previously injected with a mixture of cyp26a1 and GFP mRNA, or from control sibling embryos injected with GFP mRNA alone were cultured for 1hr (NFst10-10.25), 3hrs (NFst11), or until early neurula stage (NFst14). Microarray analysis of three independent experiments was carried out on GeneChip™ Xenopus laevis Genome 2.0 Array (Affymetrix) (Fig.1A).

**Figure 1:**
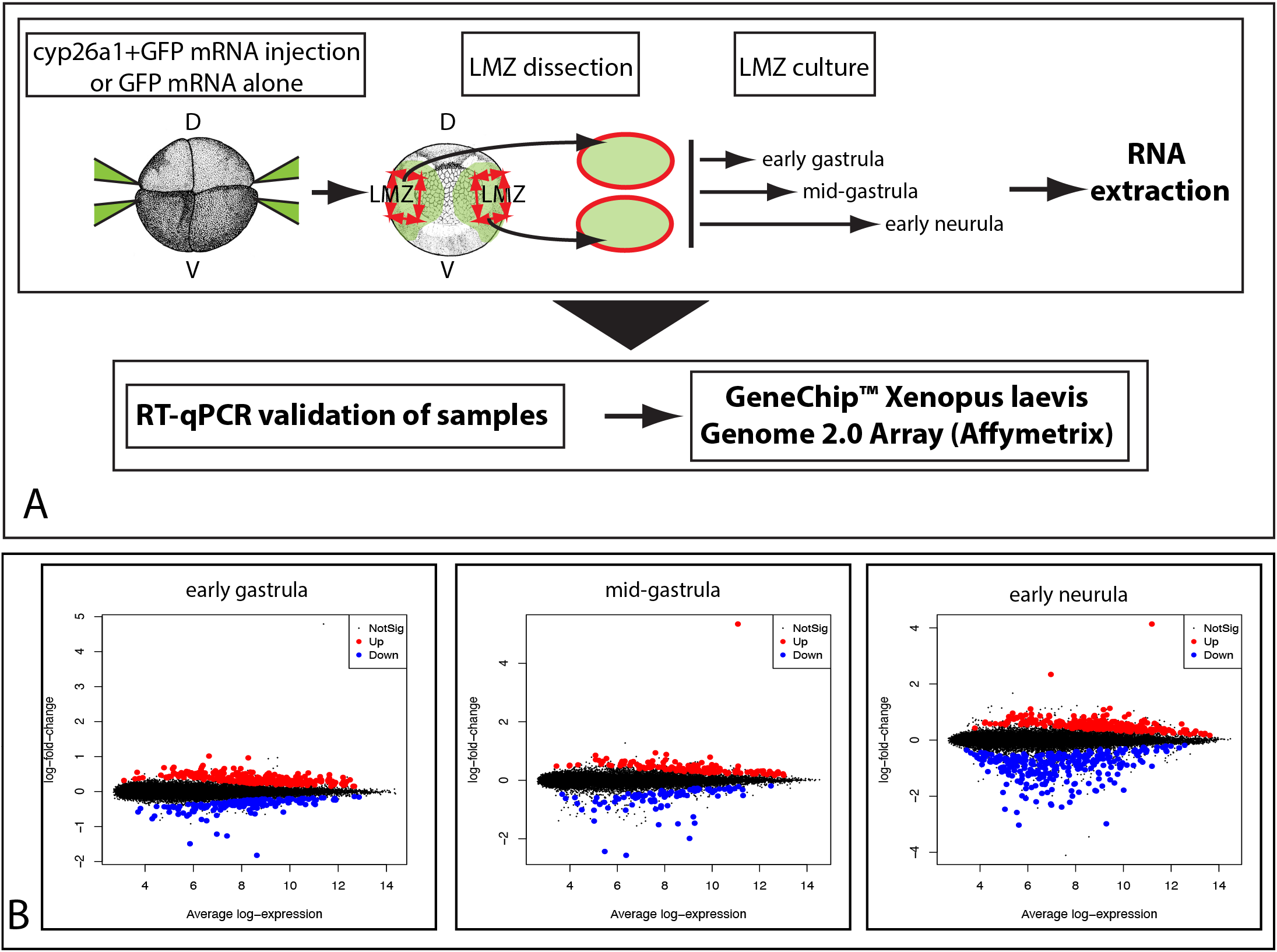
Transcriptomic profiling of RA target genes in lateral mesoderm. A. Experimental workflow. LMZ explants are dissected at early gastrula stage from embryos previously injected with cyp26a1 and GFP mRNA (disruption of RA signaling), or GFP mRNA alone (control). Profiling is performed with batches of explants cultured until early gastrula, mid-gastrula or early neurula stages. B. Overview of DE analysis. Mean-difference plots of log-intensity ratios (differences) versus log-intensity averages (means) of Affymetrix probe sets at early gastrula, mid-gastrula or early neurula stages. Black dots correspond to probe sets with fdr > 0.05, and thus excluded from DE analysis. Dots highlighted with blue correspond to probe sets whose expression is down-regulated upon RA disruption (RA-positive targets). Dots highlighted with red are probe sets whose expression is up-regulated upon RA disruption (RA-negative targets).

Differential expression (DE) analysis between control and RA-disrupted samples was performed as described in Materials and Methods. It allowed to identify genes whose expression is affected by RA disruption as soon as NFst10 (Fig1B). At this stage, 426 probe sets were differentially expressed (268 up-regulated and 158 down-regulated as a result of RA disruption) with a false discovery rate (fdr) <5% (Fig.1B). Likewise, 236 were differentially expressed at NFst11 (143 up-regulated and 93 down-regulated), and 505 at NFst14 (224 up-regulated and 281 down-regulated) (Fig.1B). Detailed results of this analysis are provided as supplementary material as “probe set analysis” tables for every stage. We further chose a log_2_fc > 0.5 (absolute value) as an arbitrary threshold for DE analysis. RA targets encoding transcription factors (TF) are listed in table 1 (positive RA targets) and supplemental table1 (negative RA targets), while a list of DE genes grouped by potential function of encoded proteins is displayed in supplemental table2. L and .S alloalleles are specified in these tables for every DE gene, but will not be further indicated in the text.

As expected, several components of the RA pathway appear as RA targets (*cyp26a, cyp26c, dhrs3, rdh13, rara, rxra*). Furthermore, *dusp6* encoding MAP kinase phosphatase-3 (MKP3) that we previously identified as RA target antagonizing FGF signaling during gastrulation (Le Bouffant et al., 2012) also appear among DE genes at Nfst10, 11 and 14, as well as *fgf3* and *fgf8* during gastrulation (Le Bouffant et al., 2012). DE at NFst14 of some identified genes was further confirmed by RT-qPCR (supplemental Fig.1A).

Because we were principally looking at transcriptional response to RA signals, we focused on probe sets corresponding to genes encoding TF down-regulated as the result of RA disruption, and thus expected to be positive RA targets (table1). At NFst14, they included TF known to play a central role in pronephros specification, like *pax8* and *lhx1* (Carroll and Vize, 1999), which have been previously shown to be down-regulated in LMZ explants by RA disruption (Le Bouffant et al., 2012), and *osr2, irx3* and *mecom* whose depletion strongly affects pronephric development (Alarcon et al., 2008; Tena et al., 2007; Van Campenhout et al., 2006). Several DE TF-encoding genes (*gbx2.2, greb1, hand1, hnf1b, mespa, mespb, nr2f2, ripply3, tbx1, tcf21, znf703*) have also been previously reported to be RA targets in various contexts (Brophy et al., 2017; Deimling and Drysdale, 2009; Gere-Becker et al., 2018; Janesick et al., 2019, 2012; Moreno et al., 2008; Tanibe et al., 2012; von Bubnoff et al., 1996). Another prominent group of DE TF corresponded to *hox* genes. Hox have also been reported to be RA targets in other processes (Nolte et al., 2019). *Hoxa1* and *d1* were detected as soon as NFst10, and thus are among the earliest RA target in LMZ. At NFst11, *hoxa1, a9, a10, b6, c5* and *d1* appeared as DE. At NFst14 DE of *hoxa2, a3, b3, b4, c5* and *d1* was detected. Although not significant in DE analysis (fdr >5%) DE of *hoxa1* at NFst14, was further confirmed by RT-qPCR, as well as that of *hoxa3, b3, b4, c5* and *d1* (supplemental Fig.1B). Finally, the hox cofactor encoding-gene *meis3* also appeared to be a prominent RA positive target of the LMZ detected at NFst11 and NFst14. This was confirmed by RT-qPCR (fig.2A). In situ hybridization (ISH) further showed that *meis3* mRNA can be detected in lateral mesoderm at neurula stage (NFst18-19), including the kidney field (Fig.2B-D).

**Figure 2:**
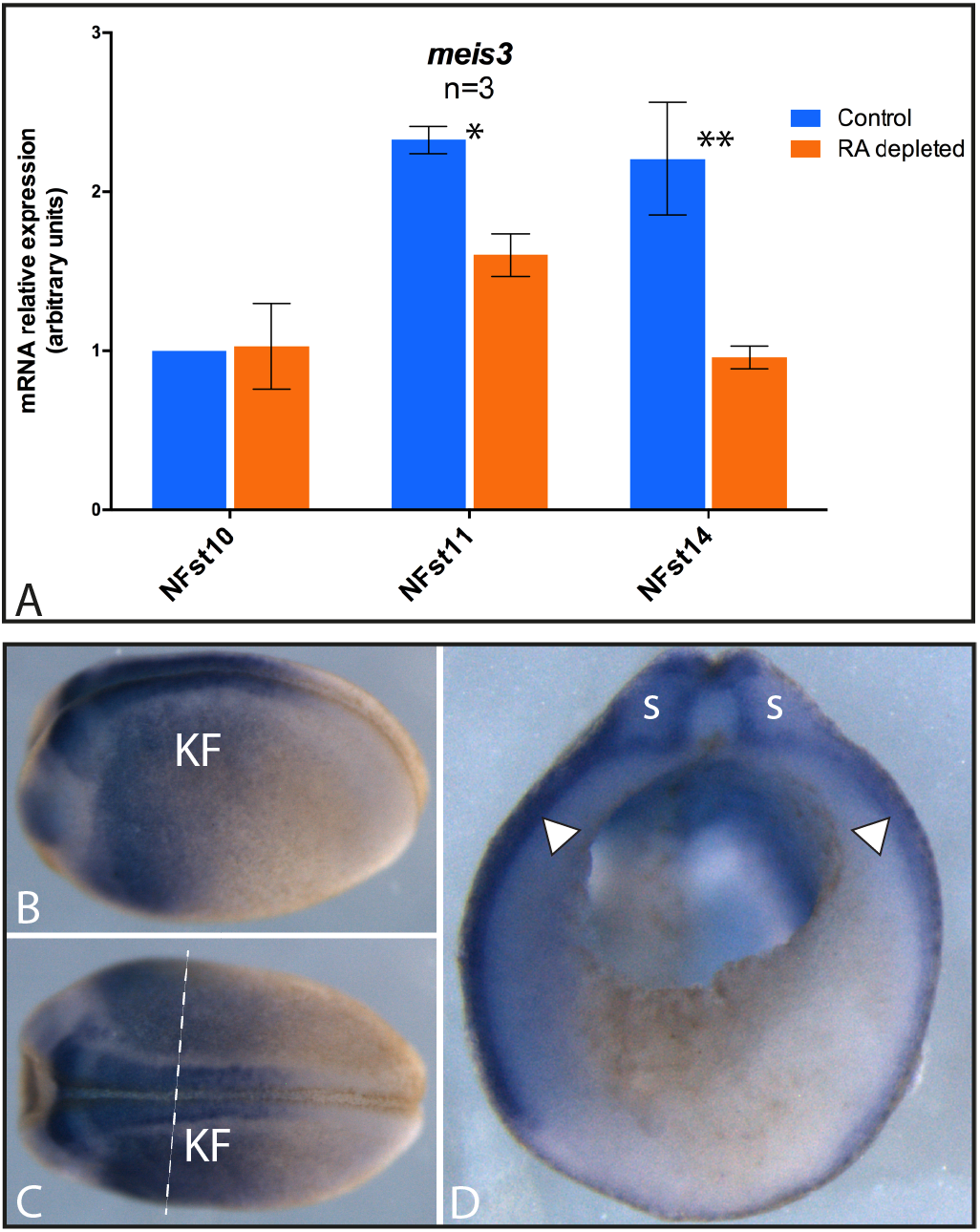
Targeted meis3 depletion results in the inhibition of *pax8* expression in the kidney field. A. RT-qPCR analysis of *meis3* expression in LMZ explants in response to RA depletion. Expression of *meis3* is shown as relative expression to *odc*. *Meis3* expression appears to be dependent on RA from midgastrula stage onward (NFst11) and at early neurula stage (NFst14). * p<0.05, ** p<0.01 (paired Student’s t-test). Error bars correspond to SEM. B-D. ISH with *meis3* antisense probe. Late neurula stage (NFst18). Lateral (B) and dorsal (C) views. D. Embryo bisected according to the dotted line shown in C. *Meis3* is expressed in dorsal and lateral mesoderm (arrowheads), including the kidney field (KF). s: somite.

### Depletion of the RA-target meis3 results in the inhibition of *pax8* expression in the kidney field

The above data raise the question of the potential role of hox downstream of RA during pronephros precursors specification. Yet, directly challenging hox function in renal precursors to evaluate their function is expected to be difficult because of the functional redundancy between the different hox expressed in these cells. We rather attempted to inhibit the hox cofactor meis3, whose expression is also regulated by RA to address this question.

We have first analyzed the effects produced by meis3 depletion upon expression of *pax8* in the kidney field. A previously characterized translation-blocking morpholino, Momeis3 (In der Rieden et al., 2011) was microinjected into the left V2 blastomere at the 8-cell stage to target ventro-lateral mesoderm at the origin of the kidney field on the left side of the embryo (Fig.3A). A 10ng dose was used that did not interfere with gastrulation (In der Rieden et al., 2011). Embryos were cultured until late neurula stage (NFst18-19), and *pax8* expression was analyzed by ISH. Comparison of *pax8* expression on left and right sides of injected embryos showed a severe inhibition of *pax8* expression in the kidney field on the left side. *Pax8* expression domain appeared much smaller, and was often fragmented into small patches (67% inhibition, n= 101, 5 independent experiments, supplemental table 3) (Fig.3B). This effect was not observed in embryos similarly injected with a control morpholino (87% normal, n= 82, 5 independent experiments, supplemental table 3) (Fig.3B). Furthermore, a mutated version of *meis3* mRNA not targeted by Momeis3 was able to significantly rescue *pax8* inhibition (Fisher’s Exact Test for Count Data p-value = 5.892e-08), when co-injected with meis3 morpholino (Fig.3C-D), demonstrating that *pax8* inhibition is a specific effect resulting from meis3 depletion and further suggesting that meis3 plays an important role in the control of *pax8* expression in the kidney field.

**Figure 3:**
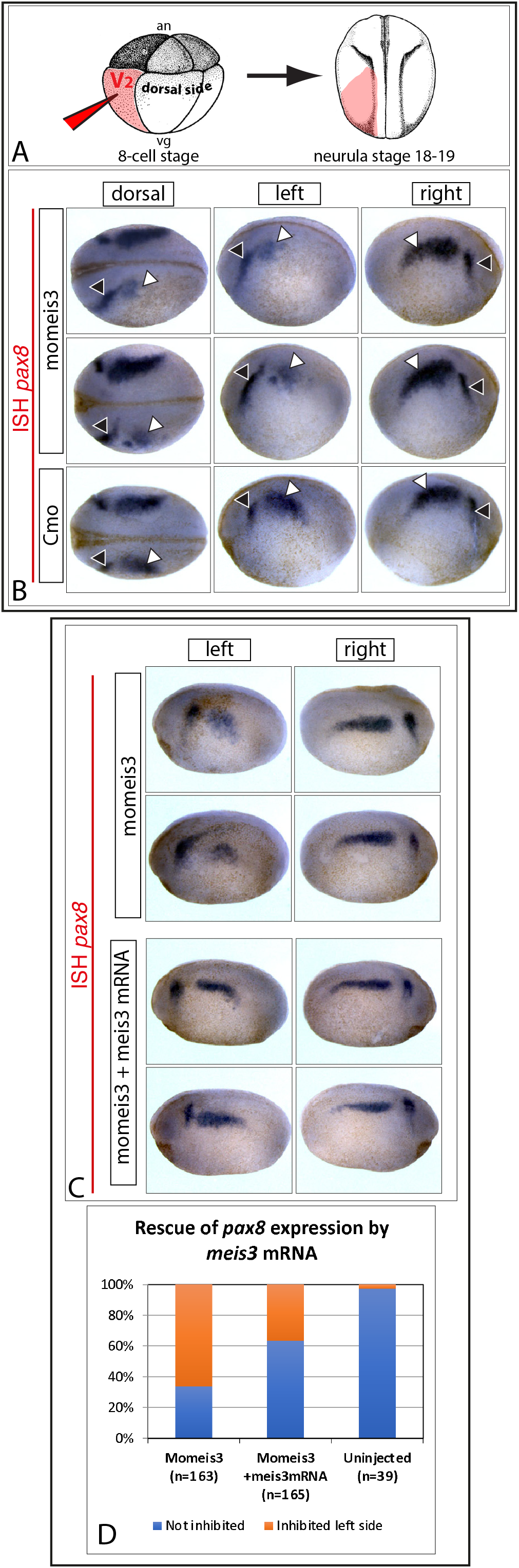
Targeted meis3 depletion results in the inhibition of *pax8* expression in the kidney field. A. Morpholinos are injected in the V2 blastomere at the 8-cell stage to target the kidney field on the left side of the embryo. Injected embryos are cultured until late neurula stage for ISH analysis of *pax8* expression. B. Representative injected embryos. Views of dorsal region, left and right sides of the same embryos are shown. White arrowheads indicate kidney field expression and black arrowhead expression in the otic placode. *Pax8* expression is strongly inhibited in the kidney field on the left side of embryos injected with Momeis3 morpholino. It is almost absent or fragmented into small patches. A control morpholino (Cmo) has no effect. C, D. Rescue experiments with a mRNA encoding a version of *meis3* transcript not targeted by Momeis3. C. Representative examples of meis3-depleted and *meis3* mRNA rescued embryos. D. Cumulative results from 3 independent experiments.

Since mesodermal expression of *aldh1a2* that encodes RA-synthesizing enzyme retinaldehyde deshydrogenase has been previously shown to involve a direct regulation by Hoxa1-Pbx1/2-Meis2 in mouse embryos, and could be down-regulated by depletion of pbx1 and hoxa1 in Xenopus embryos (Vitobello et al., 2011), a possibility is that meis3 depletion actually only interferes with RA signaling, which in turn would result in *pax8* inhibition. We have therefore analyzed to what extent meis3 depletion also affected *aldh1a2*. Meis3 depletion did not result in a strong inhibition of *aldh1a2* expression in the mesoderm similar to that observed for *pax8* (Fig.4A). *Aldh1a2* expression domain yet often appeared smaller on the injected side when compared to the control side (76% smaller, n=72, 4 independent experiments, supplemental table 3), while *aldh1a2* expression domain was mostly unaffected by control morpholino (76% unaffected, n=75, 4 independent experiments, supplemental table 3) (Fig.4A). Because *lhx1* expression in the kidney field is very sensitive to RA signaling (Cartry et al., 2006; Drews et al., 2011), we may expect a similar effect of meis3 depletion upon *lhx1*. Meis3 depletion indeed only caused a limited inhibition of *lhx1* expression. *Lhx1* expression domain appeared narrower, when compared to control side (76% narrower, n=90, 5 independent experiments, supplemental table 3) (Fig.4B), but was never affected in the same way as *pax8*. This effect was not observed in embryos injected with a control morpholino (82% unaffected, n=89, 5 independent experiments, supplemental table 3) (Fig.4B).

**Figure 4:**
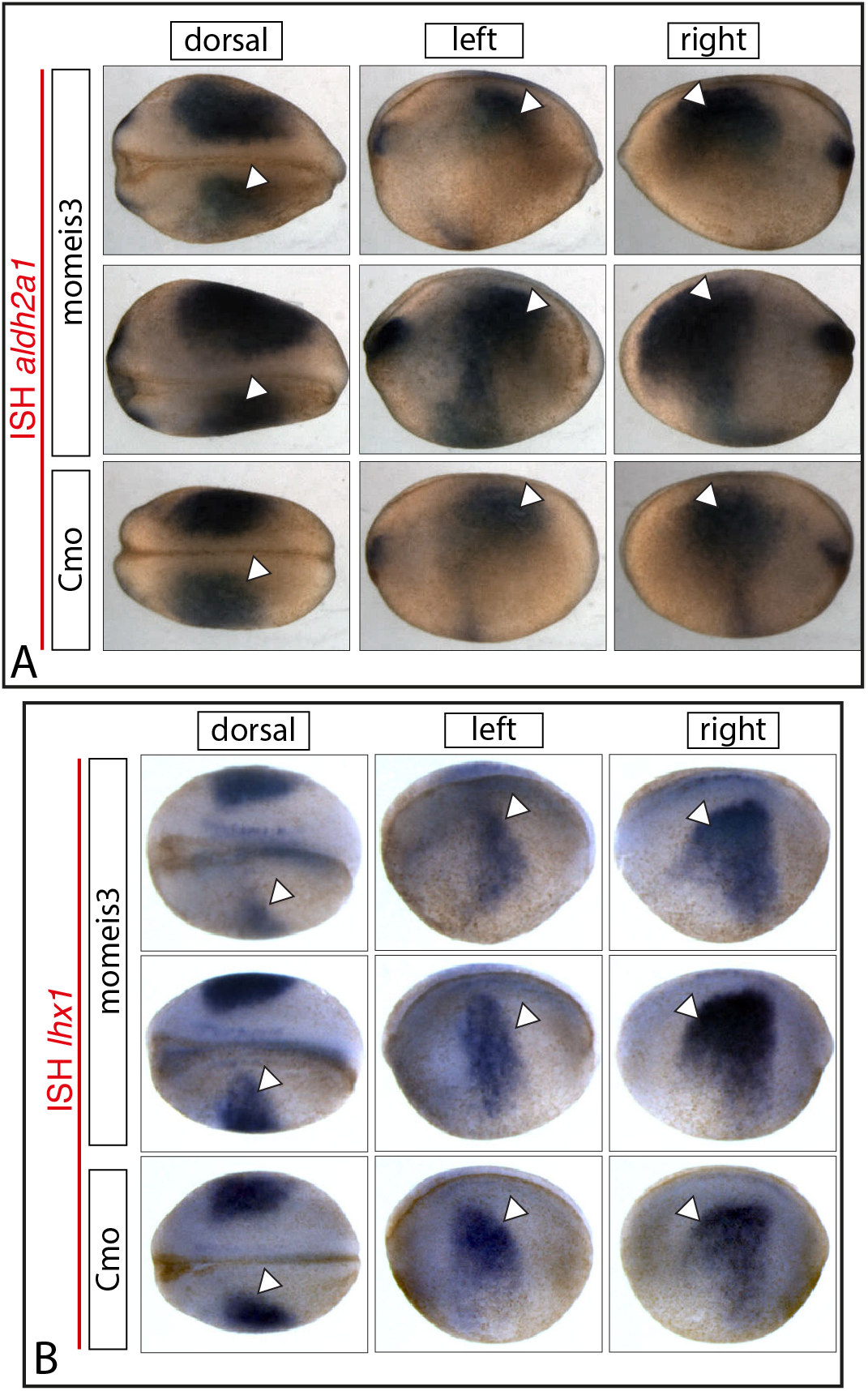
Targeted meis3 depletion has a moderate effect upon *aldh1a2* and *lhx1* expression in the kidney field. A, B. Representative injected embryos showing *aldh1a2* expression (A) or *lhx1* expression (B). A. *Adh2a1* expression is very dynamic and can vary from one embyo to the other. Yet comparing Momeis3 injected left side with control right side shows that expression domain in lateral mesoderm (arrowheads) is smaller on the injected side. This mild effect is not observed on embryos injected with a control morpholino (Cmo), although Cmo can slightly affect expression. B. *Lhx1* expression is moderately inhibited as a result of Momeis3 injection. *Lhx1* expression domain appears narrower. This is not observed after injection of Cmo, although Cmo can also slightly affect *lhx1* expression.

Together, our observations show that meis3 depletion is resulting in a strong inhibition of *pax8*, suggesting that meis3 plays an important role upstream of *pax8* expression in the kidney field. Meis3 might also act upstream of *lhx1*, but to a lower extent than for *pax8*. Meis3 might be involved in the mechanisms controlling RA synthesis in the kidney field by participating to the regulation of *aldh1a2* expression, possibly in conjunction with hoxa1 and pbx1 (Vitobello et al., 2011). Interference with RA synthesis resulting from meis3 depletion could explain the effect observed on *lhx1*. Yet, the strong effect observed on *pax8* expression is unlikely to result only from this interference, thus raising the question of other inputs acting on *pax8* that are also dependent on meis3.

### Animal cap experiments suggest an involvement of hox/pbx1 and meis3 in the control of *pax8* but not of *lhx1* expression

Since we expect meis3 to act as a hox cofactor, we further investigated whether expression in isolated blastula animal caps of meis3 associated with one hox expressed in the kidney field can cause upregulation of *pax8* and *lhx1*. Hox are often acting as hox/pbx dimers (Mann et al., 2009). We therefore tested a combination of hoxb4 that is identified as RA target (see above) and is expressed in the kidney field (In der Rieden et al., 2010), pbx1 and meis3. Messenger RNA mixtures were injected at the animal pole of every blastomere at the 4-cell stage and animal caps dissected when embryo reached the midblastula stage. Isolated caps were further cultured until sibling controls reached neurula stage NFst18-19, and were processed for RT-qPCR analysis (Fig.5A).

**Figure 5:**
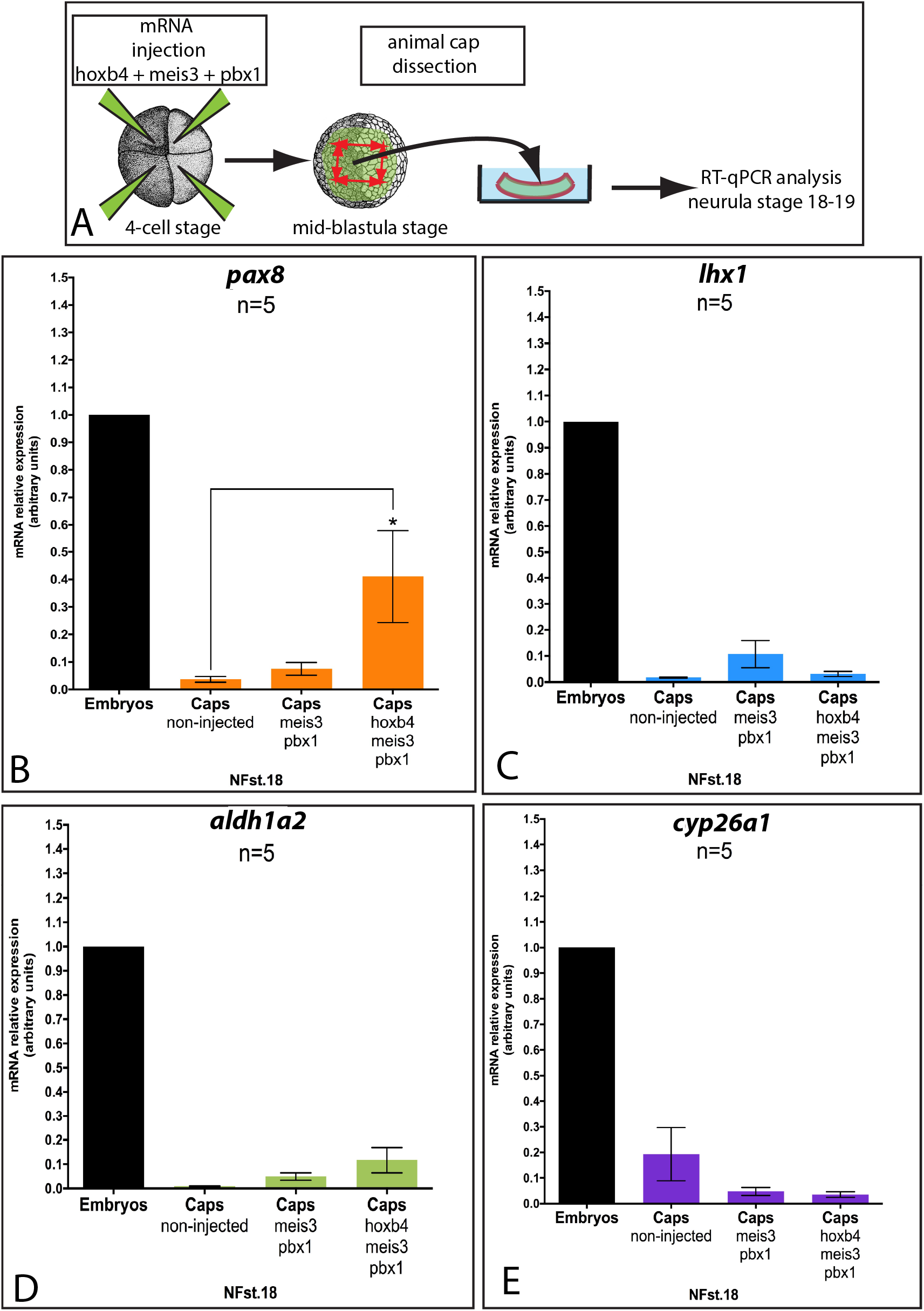
Combined expression of hoxb4, meis3 and pbx1 results in *pax8* up-regulation in animal cap assays, but not of *lhx1*. A. Mixtures of mRNA encoding Xenopus hoxb4, meis3 and pbx1 are injected into the four blastomeres of 4-cell stage embryos close to the animal pole. Animal caps are dissected at mid-blastula stage and cultured until siblings reached late neurula stage, and are processed for RT-qPCR analysis. B-E. RT-qPCR analyses. Expression of the analyzed genes is shown as relative expression to *odc*. Error bars correspond to s.e.m. B. Expression of *pax8* is up-regulated in caps expressing hoxb4 combined to meis3 and pbx1. Expression of meis3 and pbx1 alone has little effect on *pax8* expression. The same combination of hoxb4, meis3 and pbx1 neither significantly affect *lhx1* expression (C), nor RA pathway components *aldh1a2* (D) and *cyp26a1* (E). * p<0.05 (paired Student’s t-test)

The results show that hoxb4 combined with pbx1 and meis3 caused a significant up-regulation of *pax8* expression. In contrast, expression of pbx1 and meis3 alone had little effect (Fig.5B), showing that *pax8* up-regulation requires hoxb4 expression and does not result from the induction of hox expression by meis3 (In der Rieden et al., 2011). Interestingly, hoxb4 combined with pbx1 and meis3 did not cause *lhx1* up-regulation (Fig.5C), indicating that *pax8* activation does not result from a general induction of kidney field genes. We have further analyzed whether *pax8* up-regulation resulting from hoxb4 coexpressed with meis3 and pbx1 co-factors is accompanied by an effect on RA metabolism. Neither expression of *aldh1a2* (Fig.5D) nor *cyp26a1*, which encodes RA catabolizing enzyme cyp26 was significantly affected by the expression of hoxb4, meis3 and pbx1 (Fig.5D, E). These experiments show that combining a hox with pbx1 and meis3 cofactor can upregulate *pax8* in a process that does not rely on RA synthesis, raising the question of a direct input of hox and hox cofactors on *pax8* expression.

### Transactivation of Pax8-CNS1 by *Xenopus* hoxb4, meis3 and pbx1 suggests a direct regulation of *pax8* by hox

*Pax8* regulation in the late pronephric anlage at tailbud stage involves a set of conserved enhancers and a silencer located in the proximal promoter region of the gene that restricts *pax8* expression to the pronephros. Four conserved enhancers (Pax8-CNS1-4) can drive reporter expression in the pronephric anlage in transgenics at tailbud stages when associated to a minimal promoter. One of these conserved enhancers, Pax8-CNS1 contains three conserved putative Pbx-Hox binding motifs and one putative Meis motif (Fig.6A). When it is associated with the silencer region, expression is restricted to the pronephros in transgenesis assays (Ochi et al., 2012). These data raise the question of a regulation of this enhancer by hox. We have therefore tested whether various combinations of Xenopus hoxb4, pbx1 and meis3 can transactivate Pax8-CNS1 in HEK293 cells.

**Figure 6:**
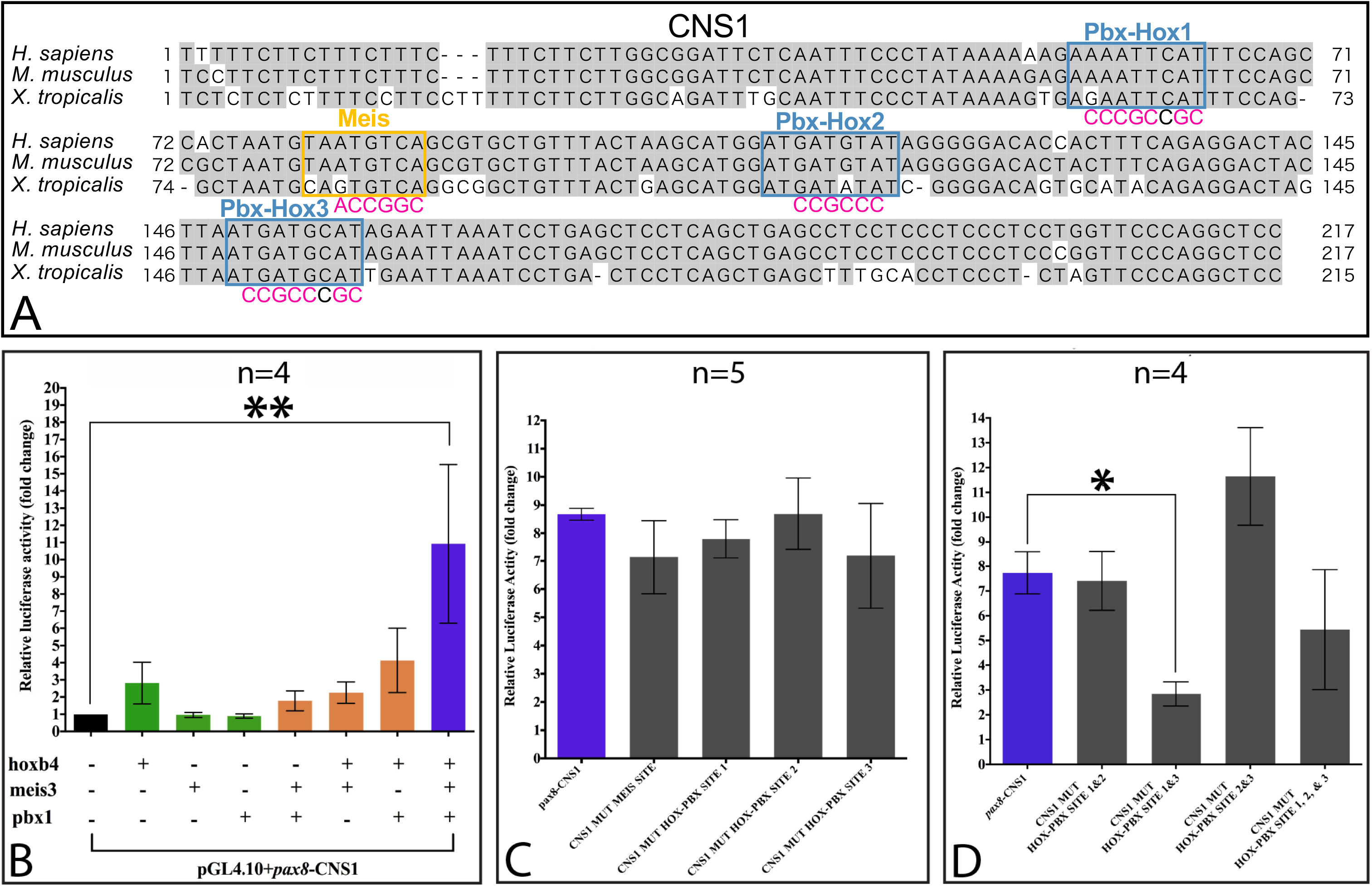
Transactivation of Pax8-CNS1 by Xenopus hoxb4 combined with meis3 and pbx1. A. Alignment of *pax8*-CNS1 sequences from the human (hg38, chr2: 113,341,953-113,342,169), mouse (mm10, chr2: 24,539,076-24,539,292), and frog (*Xenopus tropicalis*; xenTro10, chr3: 125,792,358-125,792,572) with three highlighted conserved putative Pbx-Hox binding motives (blue), and one Meis (orange). Nucleotides identical in two or three sequences are shaded in grey. Mutations tested for every site are shown in red. Note that the putative binding motives mapped here show close similarity to the consensus binding sequences of Pbx-Hox (5’-ATGATTNATNN-3’) or Meis (5’-TGACAGST-3’) (Chang et al., 1997, 1996). B. Reporter expression measured for HEK293 cells transfected with pax8CNS1-luc, and various combinations of plasmids allowing expression of Xenopus hoxb4, meis3 or pbx1 indicated in the bottom of the figure. Values are expressed as fold changes relative to HEK293 cells transfected with pax8CNS1-luc alone. A significant increase of reporter expression is only observed with cells expressing a combination of hoxb4, meis3 and pbx1. Other combinations tested do not elicit any significant increase of reporter expression. ** p<0.01Dunn’s multiple comparison test after Kruskal-Wallis test). C. Analysis of mutations of single Pbx-Hox and Meis motifs in transactivation assays. HEK293 cells were transfected with the three plasmids allowing expression of hoxb4, meis3 and pbx1 combined with wild-type pax8CNS1-luc, pax8CNS1mut-lucMeis, pax8CNS1mut-lucHP1, pax8CNS1mut-lucHP2 or pax8CNS1mut-lucHP3. Values are expressed as fold changes relative to HEK293 cells transfected with wild-type pax8CNS1-luc. None of the single mutation tested significantly affected transactivation. D. Analysis of multiple mutations of Pbx-Hox motifs in transactivation assays. HEK293 cells were transfected with the three plasmids allowing expression of hoxb4, meis3 and pbx1 combined with wildtype pax8CNS1-luc, pax8CNS1mut-lucHP1+2, pax8CNS1mut-lucHP1+3, pax8CNS1mut-lucHP2+3 or pax8CNS1mut-lucHP1+2+3. Values are expressed as fold changes relative to HEK293 cells transfected with wild-type pax8CNS1-luc. Combining mutation of Pbx-Hox 1 and 3 motifs results in a strong decrease of reporter expression, pointing to these two sites as active hox-Pbx binding sites. * p<0.05 (Dunn’s multiple comparison test after Kruskal-Wallis test).

HEK293 cells were transfected with a reporter construct containing Pax8-CNS1 sequence upstream of a minimal promoter followed by the luciferase coding sequence. HEK293 cells were either transfected with the reporter construct alone, or were co-transfected with the reporter construct and vectors allowing expression of the various combinations of transcription factors tested (see Materials and methods). Combination of hoxb4, pbx1 and meis3 elicited a reporter activation 10 fold higher than the basal activation observed in cells transfected with the reporter construct alone (Fig6B) (4 independent experiments, p<0.01, Dunn’s multiple comparison test after Kruskal-Wallis test). This strong effect was never observed when cells were co-transfected with hoxb4, meis3 or pbx1 alone, or combinations of pbx1 and meis3, hoxb4 and meis3, or hoxb4 and pbx1 (Fig.7B), suggesting that the three transcription factors are cooperating.

**Figure 7:**
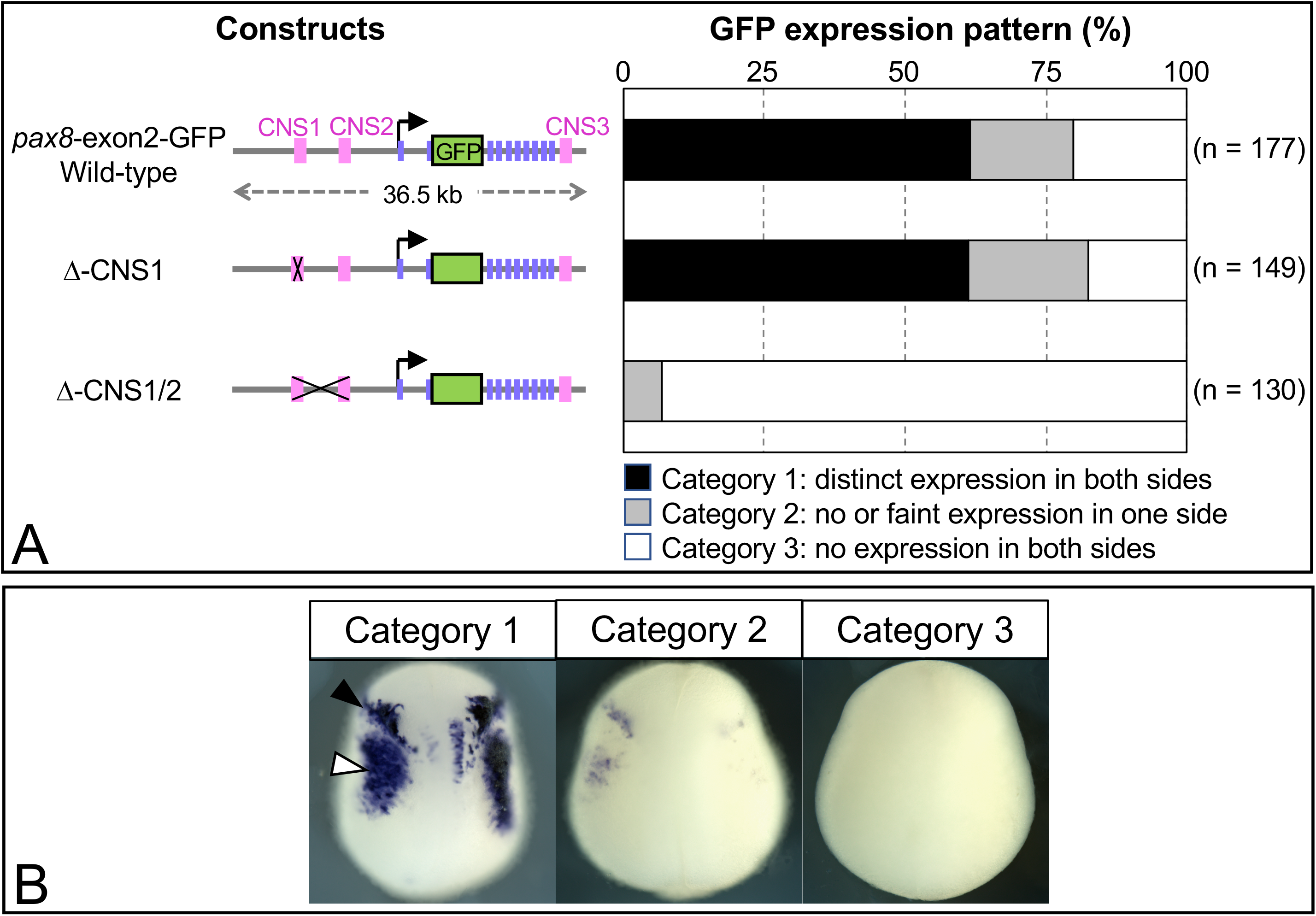
A genomic region encompassing the CNS1 and CNS2 is required for otic placode- and pronephric kidney field-specific expression of *pax8*. A. GFP reporter constructs used for transgenesis experiments in *X. laevis* embryos (left) and bar graphs summarizing GFP expression patterns in the resulting embryos (right). The bar graphs show ratio of embryos showing distinct otic placode- and kidney field-specific expression in both sides (scored as the Category 1), no or faint expression in one side (Category 2), and no expression in both sides (Category 3). B. Representative embryos showing GFP expression scored as the Category 1, Category 2, and Category 3; dorsal view with anterior side to the top. Black and white triangles indicate expression in the otic placode and kidney field, respectively.

In order to identify which putative binding sites are involved during transactivation, reporter construct bearing different combinations of mutations of any of the three Pbx-Hox motifs, or of the Meis motif (Fig.6A), were tested in transactivation assays. Neither single mutations of Pbx-Hox motifs nor mutation of the Meis motif appeared to affect reporter activation (Fig.6C). The putative Meis motif present in Pax8-CNS1 is therefore not required for transactivation, since similar mutations of Meis binding sites have been previously shown to interfere with Meis2 interaction (Wassef et al., 2008). This suggests that either meis3 is interacting with another site of Pax8-CNS1, or that meis3 does not need to bind to DNA to exert its effect. The absence of effect of single mutations of any of the Pbx-Hox binding motives may rather suggest redundancy. We therefore tested combinations of PBX-Hox sites mutations. Combining mutations of Pbx-Hox motif 1 and 3 resulted in a strong inhibition of reporter activation (Fig.6D) (4 independent experiments, p<0.05, Dunn’s multiple comparison test after Kruskal-Wallis test), showing that these two sites are required for transactivation. In contrast, combining mutation of Pbx-Hox motif 2 with mutations of Pbx-Hox motives 1 or 3 does not have any significant effect on reporter activation, suggesting that this motif is not used. Together these results support the idea of a potential direct regulation of *pax8* by hox.

### A region encompassing Pax8-CNS1 and Pax8-CNS2 is critical for *pax8* regulation at neurula stage

In order to identify regulatory sequences involved in *pax8* regulation in the pronephric kidney field, we have performed reporter assays using a previously characterized transgene encompassing 36.5 kb of the *X. tropicalis pax8* gene sequence with a GFP cassette inserted into the second exon. This transgene allows expression of a truncated pax8-GFP fusion protein. Transgenesis experiments in *X. laevis* embryos with this sequence have shown that expression of this transgene recapitulates endogenous *pax8* expression in the developing pronephros and otic vesicle (Ochi et al., 2012). The 36.5kb sequence includes three of the conserved *pax8* enhancers, Pax8-CNS1, Pax8-CNS2 and Pax8-CNS3 (Fig.7A). We have first tested whether removing Pax8-CNS1 from the transgene affected its expression. Reporter expression was monitored by ISH at late neurula stage (NFst18/19). Transgenesis experiments performed with the wild-type transgene, or with the transgene lacking Pax8-CNS1 (Δ-CNS1) showed that the absence of Pax8-CNS1 did not affect transgene expression; more than 61% of the embryos manipulated with the wild-type or the Δ-CNS1 construct showed otic placode- and kidney field-specific GFP expression in both sides of their bodies (n = 109/177 and n = 91/149, respectively; Fig.7A, B). It is possible that the presence of Pax8-CNS2 and/or Pax8-CNS3 may compensate for the absence of Pax8-CNS1. Previous transgenesis analyses performed with these different sequences associated to a minimal promoter have indeed shown that they all drive reporter expression in the pronephric anlage at tailbud stages (Ochi et al., 2012). We have therefore tested whether removing Pax8-CNS1 and Pax8-CNS2 from the transgene would affect its expression. Transgenesis experiments were carried out with a transgene deleted of a 3.5kb sequence encompassing Pax8-CNS1 and Pax8-CNS2 (Δ-CNS1/2, Fig.7A). This resulted in a dramatic inhibition of reporter expression: none of the manipulated embryos showed GFP expression in both sides and less than 7% of them showed otic- and kidney field-specific GFP expression only in one side of their bodies (n = 9/130, Fig.7A, B). These results point to this 3.5kb region as critical for *pax8* regulation both in kidney field and the otic placode. It contains two conserved enhancers Pax8-CNS1 and Pax8-CNS2, and could be regulated by hox at the level of Pax8-CNS1.

## Discussion

RA signaling is centrally involved in the specification of pronephric precursors of the kidney field by controlling the expression of master transcriptional regulators of renal development such as pax8 and lhx1 in a subset of lateral mesodermal cells (Cartry et al., 2006; Drews et al., 2011). The present study provides evidence for a role of hox in the control of *pax8* expression downstream of RA. Expression profiling of RA target genes in lateral mesendoderm during gastrulation and early neurulation allowed to identify several hox genes, including *hoxa1, a3, b3, b4, c5* and *d1* as RA targets, as well as *meis3* that encodes a TALE homeodomain hox cofactor. We further show that targeted depletion of meis3 in the kidney field results in an inhibition of *pax8* expression. *Pax8* can also be induced in response to hoxb4 combined with cofactors meis3 and pbx1 in ectodermal animal caps. Supporting the view of a direct control of *pax8* expression by hox, the same combination of TF is able to transactivate a previously identified pax8 enhancer, Pax8-CNS1 (Ochi et al., 2012),. Using transgenesis experiments with reporter fosmid constructs containing −24kb to +12kb of the Xenopus *pax8* gene that recapitulate endogenous *pax8* expression, we further show that deletion of a 3.5kb region encompassing Pax8-CNS1 and a second *pax8* enhancer, Pax8-CNS2, totally abolishes reporter expression in the kidney field, showing that this region plays a critical role in *pax8* regulation. Deletion of Pax8-CNS1 alone is yet not sufficient to affect reporter expression, indicating that hox may also interact with other regions of this 3.5kb sequence, and/or that other inputs are cooperating with hox during regulation of *pax8* in renal precursors of the kidney field.

Hox are thought to play an important role in the control of RA synthesis in paraxial and lateral mesoderm through the control of *aldh1a2* expression. Expression of *aldh1a2* in paraxial and lateral mesoderm of Xenopus embryos has been indeed previously shown to be inhibited by depletion of hoxa1 and pbx1. Direct regulation of mouse Aldh1a2 was further shown to involve the interaction of hoxa1-pbx1/2-meis2 complex with a regulatory element that is required to maintain normal *Aldh1a2* expression levels in mouse embryos, and a mechanism maintaining high levels of RA in mesoderm through a feed-forward loop has been proposed (Vitobello et al., 2011). The present results show that meis3 depletion only marginally affects *aldh1a2* expression, while it strongly affects *pax8*. This suggests that besides controlling RA synthesis, hox might also regulate *pax8* in a more direct way. Although in an ectodermal context, our observations showing that a combination of hoxb4, pbx1 and meis3 is able to up-regulate *pax8* in animal caps without significantly inducing *aldh1a2* expression further support this view. Finally, transactivation of Pax8-CNS1 by hox and cofactors suggests a direct regulation in this process.

Although the present study points to hox in the control of *pax8* downstream of RA, it is likely that *pax8* expression in the kidney field does not depend only on hox, and hox may cooperate with other RA targets, as summarized in Fig.8. *Lhx1* can be a direct target of RA, since it is up-regulated in the embryo and in animal caps as an immediate-early response to exogenous RA (Cartry et al., 2006; Drews et al., 2011). Depletion of lhx1 results in a reduction of the *pax8* expression domain of the kidney field, while expression of constitutively active forms of lhx1 expands this domain (Cirio et al., 2011), suggesting that lhx1 may also act upstream of pax8 in renal precursors. A direct regulation of *pax8* by lhx1 still remains elusive. RA can also influence *pax8* expression in a more indirect way. We have previously shown that TRPP2-dependent intracellular calcium signaling is required for *pax8* expression in the kidney field. Both intracellular calcium signaling and expression of the TRPP2 channel at the plasma membrane were inhibited upon RA disruption (Futel et al., 2015). It is not known whether this increase in cytoplasmic calcium transients also results in an intranuclear calcium increase that could directly influence *pax8* transcriptional control. Finally, another RA target *dusp6* that encodes the enzyme mkp3 can also positively influence *pax8* expression in the kidney field by inhibiting the FGF/erk pathway known to impede early pronephric development (Colas et al., 2008; Le Bouffant et al., 2012).

**Figure 8:**
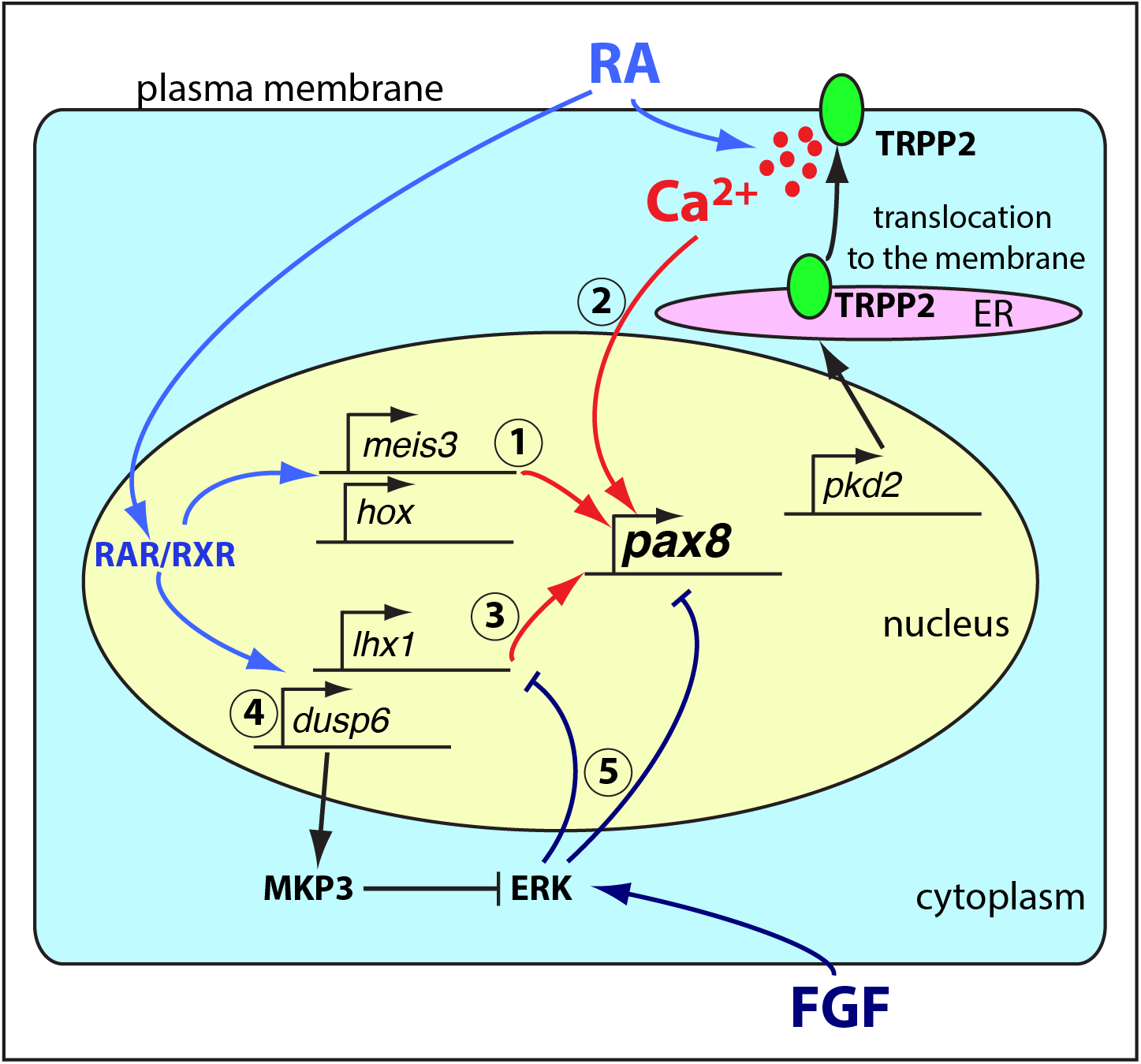
Working model based on present data and previously published results (Cartry et al., 2006; Cirio et al., 2011; Colas et al., 2008; Futel et al., 2015; Le Bouffant et al., 2012) illustrating the mechanisms that control *pax8* expression in the kidney field. The present data (1) show that meis3 and hox are RA targets, and are involved in the direct regulation of *pax8*. They are cooperating with other RA-dependent inputs such as lhx1 (Cirio et al., 2011) (3), and a yet unidentified mechanism dependent on RA-dependent intracellular calcium signaling (Futel et al., 2015) (2). The FGF/erk pathway was previously shown to inhibit *pax8* expression (Colas et al., 2008) (5) and can be negatively regulated by MKP3 that is encoded by *dups6*, another RA target (Le Bouffant et al., 2012)(this study) (4). ER: endoplasmic reticulum.

We have identified a 3.5kb regulatory region of the *pax8* gene that is absolutely required for its expression at neurula stage. This region includes two previously identified enhancers, Pax8-CNS1 and Pax8-CNS2 (Ochi et al., 2012). Reporter experiments show that a construct lacking Pax8-CNS1 is still able to drive reporter expression both in otic placode and kidney field expression domains of *pax8*. This suggests that additional inputs are interacting with other regions of the 3.5kb sequence, especially at the level of Pax8-CNS2. It is possible that this enhancer can be also regulated by hox. It indeed contains two conserved Hox-Pbx binding motives and one Meis motif, and bound by Hox and Meis proteins in a human acute lymphoblastic leukemia cell line (supplemental figure 2). As outlined above, it is also likely that TF(s) different from hox and their cofactors may be also interacting with this 3.5kb sequence. One candidate is lhx1 and it would be interesting to test if the constitutively active ldb1-lhx1 protein can transactivate Pax8-CNS2, or the whole 3.5kb sequence. As outlined above, intracellular calcium signaling might also be involved in the transcriptional control of pax8. For example, calcium-dependent release of Pax8 repression by DREAM-kcnip3 proteins has been reported in rat thyroid cells (D’andrea et al., 2005). However, DREAM-kcnip3 repression would be expected to occur close to the transcription start site (D’andrea et al., 2005), which is not included in the 3.5kb sequence that has been identified here. Cytosolic calcium can also influence pax8 expression more indirectly through more complex calmoduline kinase-dependent phorphorylation processes as those described in cardiomyocytes (Dewenter et al., 2017).

Our results show that hoxb4 both causes upregulation of *pax8* in animal caps and transactivation of Pax8-CNS1 when it is associated with pbx1 and meis3. *Hoxb4* is expressed in lateral mesoderm, but several other hox genes have been identified as RA targets in this study, and are likely expressed in lateral mesoderm where they can also be potentially involved in *pax8* regulation. Complexity of *hox* gene expression in pronephric precursors is in line with the even higher complexity of *hox* gene expression observed during mouse metanephric kidney development (Patterson and Potter, 2004). *Hox* genes often display a high degree of overlapping expression and functional redundancy that have made functional analyses very challenging. For example, knockout of the four *hox* PG9 genes does not lead to kidney defect (Xu and Wellik, 2011), although these genes are all expressed in the developing kidney, including renal vesicle, branching tubule, proximal, intermediate and distal segments of S-shaped bodies (Patterson and Potter, 2004). Triple knockout of *hox* PG10 genes only affects cortical cell differentiation and integration (Yallowitz et al., 2011). Important functions of hox genes have been yet identified in the developing mouse kidney. Triple knockout of *hox* PG11 genes results in kidney agenesis (Wellik et al., 2002). Hox11 proteins have been shown to form a complex with Eya1 and Pax2 proteins to regulate expression of *Six2* and *Gdnf* genes (Gong et al., 2007). Deletions of multiple loci in the hoxd cluster also revealed a role in the control of apoptosis with formation of cystic kidneys (Di-Poi et al., 2007). Hox genes are also probably playing important functions in the acquisition of nephron cell type identities. For example, cells expressing proximal tubule markers differentiate in the vicinity of cells expressing distal tubule markers in *Hoxc9,10,11−/− Hoxd9,10,11−/−* mutants. The mechanisms involved are poorly understood, but appear to be distinct from those governing acquisition of tubule segment identity (Drake et al., 2018). Functional studies carried out during mouse kidney development have not revealed a role of *hox* genes in the control of either *Pax2* or *Pax8* expression. It is unclear, however, whether a role for hox in the regulation of *pax8* is restricted to pronephric precursors, or if a similar function of hox genes could be uncovered by analyzing other combinations of *hox* gene knockouts during mouse kidney development.

In summary, we have shown that hox and their cofactor meis3 are transcriptional targets of RA in lateral mesoderm at the origin of the pronephric kidney field. Our results further show that meis3 is playing a central role in the control of *pax8* expression in the developing renal precursors during neurulation and can directly regulate pax8 expression in association with hox/pbx. Future work will identify additional inputs cooperating with meis3 and hox/pbx to finely tune *pax8* expression in developing renal precursors.

## Aknowledgements

We thank S. Authier, A. Bois and E. Manzoni for excellent technical assistance in the maintenance of the Xenopus animal facility. We are also extremely grateful to D. Gentien at Genomics platform at Curie Institute for Affymetrix microarray analysis, as well as to C. Antoniewski at the ArtBio bioinformatics platform of IBPS for transcriptomics data interpretation. We also thank Dr AH Monsoro for the *X. laevis* meis3 clone.

## Supplemental data

Probe_set_analysis_NFst10.xlsx

Probe_set_analysis_NFst11.xlsx

Probe_set_analysis_NFst14.xlsx

Supplemental table 1.docx

Supplemental table 2.docx

Supplemental table 3.docx

Supplemental table 4.docx

Legend to supplementary figure 1:

RT-qPCR validation of RA targets. A. Effect of RA depletion upon various RA targets identified by expression profiling. Expression is shown as relative expression to *β-actin*. B. Effect of RA depletion upon hox gene expression. Expression is shown as relative expression to *odc (hoxa3, b3, b4*) or *β-actin (hoxa1, c5, d1*). * p<0.05, ** p<0.01 (paired Student’s t-test). Error bars correspond to SEM.

Legend to supplementary figure 2:

*pax8*-CNS2 may be regulated by Hox and Meis proteins. A. Alignment of *pax8*-CNS2 sequences from the human (hg38, chr2: 113,324,980-113,325,232), mouse (mm10, chr2: 24,521,500-24,521,747), and frog (*Xenopus tropicalis*; xenTro10, chr3: 125,795,651-125,795,890) with two highlighted conserved putative Pbx-Hox binding motives (blue), and one Meis (orange). These putative binding motives show close similarity to the consensus binding sequences of Pbx-Hox (5’-ATGATTNATNN-3’) or Meis (5’-TGACAGST-3’). Nucleotides identical in two or three sequences are shaded in grey. B. ChIP-seq analysis indicates that MEIS1 and HOXA9 bind to human CNS2 in an acute lymphoblastic leukemia cell line (SEM). The ChIP-seq mapping data collected in ReMap2020 database (Hammal et al., 2022) was shown with schematic Multiz alignments of the human, mouse and *X. tropicalis* CNS2 sequences on UCSC Genome Browser (Kent et al., 2002).

