## Supplemental Material for "Retinoic acid control of pax8 during renal specification of Xenopus pronephros involves hox and meis3" (from another version of this preprint: version 1, 2022-06-21)

### Supplemental table 1

| Genes | Xenbase.gene.ID | Description | St10 | St11 | St14 |
|---|---|---|---|---|---|
| atf3.L | XB-GENE-6085256 | activating transcription factor 3 [Source:Xenbase;Acc:XB-GENE-6085251] | + | - | - |
| cited4.L | XB-GENE-946644 | Cbp/p300-interacting transactivator [Source:Xenbase;Acc:XB-GENE-946639] | - | - | + |
| eomes.L | XB-GENE-865305 | eomesodermin [Source:Xenbase;Acc:XB-GENE-479769] | - | - | + |
| foxa1.L | XB-GENE-487356 | forkhead box A1 [Source:Xenbase;Acc:XB-GENE-487352] | - | + | - |
| foxa2.L | XB-GENE-865381 | forkhead box A2 [Source:Xenbase;Acc:XB-GENE-480476] | + | - | + |
| foxd4l1.1.S | XB-GENE-865254 | forkhead box D4 like 1, gene 1 [Source:Xenbase;Acc:XB-GENE-479247] | - | - | + |
| hesx1.S | XB-GENE-6252654 | HESX homeobox 1 [Source:Xenbase;Acc:XB-GENE-483193] | + | - | - |
| irf1.L | XB-GENE-6255163 | interferon regulatory factor 1 [Source:Xenbase;Acc:XB-GENE-942994] | + | - | - |
| irf1.S | XB-GENE-17345840 | interferon regulatory factor 1 [Source:Xenbase;Acc:XB-GENE-942994] | + | - | - |
| irf6.2.L | XB-GENE-5910427 | interferon regulatory factor 6, gene 2 [Source:Xenbase;Acc:XB-GENE-5910331] | - | - | + |
| jun.L | XB-GENE-864843 | jun proto-oncogene [Source:Xenbase;Acc:XB-GENE-482228] | + | - | - |
| klf10.S | XB-GENE-866571 | Kruppel-like factor 10 [Source:Xenbase;Acc:XB-GENE-854761] | - | + | - |
| ncoa2.S | XB-GENE-864856 | nuclear receptor coactivator 2 [Source:Xenbase;Acc:XB-GENE-482311] | - | + | - |
| nfkbia.L | XB-GENE-866028 | NFKB inhibitor alpha [Source:Xenbase;Acc:XB-GENE-491079] | - | + | - |
| nfkbia.S | XB-GENE-17331094 | NFKB inhibitor alpha [Source:Xenbase;Acc:XB-GENE-491079] | - | + | - |
| phox2a.S | XB-GENE-853858 | paired like homeobox 2A [Source:Xenbase;Acc:XB-GENE-853857] | - | - | + |
| rreb1.L | XB-GENE-6488261 | ras responsive element binding protein 1 [Source:Xenbase;Acc:XB-GENE-492150] | - | - | + |
| rxra.L | XB-GENE-865207 | retinoid X receptor alpha [Source:Xenbase;Acc:XB-GENE-478718] | - | - | + |
| sox17b.1.L | XB-GENE-865510 | SRY (sex determining region Y)-box 17 beta, gene 1 [Source:Xenbase;Acc:XB-GENE-484865] | - | - | + |
| sox7.L | XB-GENE-865761 | SRY-box 7 [Source:Xenbase;Acc:XB-GENE-488067] | - | - | + |
| sp5.L | XB-GENE-6254266 | Sp5 transcription factor [Source:Xenbase;Acc:XB-GENE-1217606] | - | - | + |
| srebf2.L | XB-GENE-1011364 | sterol regulatory element binding transcription factor 2 [Source:Xenbase;Acc:XB-GENE-1011360] | - | - | + |
| tbx2.S | XB-GENE-6255176 | T-box 2 [Source:Xenbase;Acc:XB-GENE-1018115] | - | - | + |
| vegt.S | XB-GENE-1033836 | vegt protein [Source:Xenbase;Acc:XB-GENE-1033829] | - | - | + |
| zdhhc2.L | XB-GENE-947261 | zinc finger DHHC-type containing 2 [Source:Xenbase;Acc:XB-GENE-947256] | - | - | + |
| znf217.L | XB-GENE-1020935 | zinc finger protein 217 [Source:Xenbase;Acc:XB-GENE-1020928] | - | - | + |
| znf750.L | XB-GENE-5775750 | zinc finger protein 750 [Source:Xenbase;Acc:XB-GENE-5775684] | - | - | + |

Supplemental table1 : Retinoic acid negative targets encoding transcription factors (genes up-regulated as a result of retinoic acid disruption with a $log_2fc \geq 0.5$). Manual annotation of some genes from Xenbase website pages information is indicated in red.

### Supplemental table 2

| RA pathway | | | | | |
|---|---|---|---|---|---|
| Genes | Xenbase.gene.ID | description | St 10 | St 11 | St 14 |
| cyp26a1.L | XB-GENE-866599 | cytochrome P450 family 26 subfamily A member 1 [Source:Xenbase;Acc:XB-GENE-855583] | + | + | + |
| cyp26c1.S | XB-GENE-1219119 | cytochrome P450 family 26 subfamily C member 1 [Source:Xenbase;Acc:XB-GENE-1219115] | - | - | + |
| dhrs3.L | XB-GENE-955331 | dehydrogenase/reductase (SDR family) member 3 [Source:Xenbase;Acc:XB-GENE-955324] | + | + | + |
| rara.L | XB-GENE-865415 | retinoic acid receptor alpha [Source:Xenbase;Acc:XB-GENE-480836] | - | - | + |
| cyp26a1.S | XB-GENE-17332184 | cytochrome P450 family 26 subfamily A member 1 [Source:Xenbase;Acc:XB-GENE-855583] | - | + | + |
| rdh13.L | XB-GENE-965073 | retinol dehydrogenase 13 (all-trans/9-cis) [Source:Xenbase;Acc:XB-GENE-965068] | + | - | - |
| FGF pathway | | | | | |
| Genes | Xenbase.gene.ID | description | St 10 | St 11 | St 14 |
| dusp6.S | XB-GENE-17333168 | dual specificity phosphatase 6 [Source:Xenbase;Acc:XB-GENE-978216] | + | + | + |
| fgf2.L | XB-GENE-487006 | fibroblast growth factor 2 [Source:Xenbase;Acc:XB-GENE-487002] | - | - | + |
| fgf3.S | XB-GENE-1018521 | fibroblast growth factor 3 [Source:NCBI gene;Acc:493516] | - | + | - |
| fgf8.L | XB-GENE-865429 | fibroblast growth factor 8 [Source:Xenbase;Acc:XB-GENE-480983] | + | - | - |
| fgfr3.L | XB-GENE-479259 | fibroblast growth factor receptor 3 (achondroplasia, thanatophoric dwarfism) [Source:Xenbase;Acc:XB-GENE-479255] | - | - | + |
| fgfrl1.L | XB-GENE-482476 | fibroblast growth factor receptor like 1 [Source:Xenbase;Acc:XB-GENE-482472] | - | - | + |
| Wnt pathway | | | | | |
| Genes | Xenbase.gene.ID | description | St 10 | St 11 | St 14 |
| fzd8.S | XB-GENE-6254029 | frizzled class receptor 8 [Source:Xenbase;Acc:XB-GENE-480800] | - | + | - |
| sfrp2.S | XB-GENE-485151 | secreted frizzled-related protein 2 [Source:Xenbase;Acc:XB-GENE-485147] | - | - | + |
| trabd2a.S | XB-GENE-6488536 | TraB domain containing 2A - Metalloprotease that acts as a negative regulator of the Wnt pathway [Source:Xenbase;Acc:XB-GENE-6050964] | - | - | + |
| wnt11b.L | XB-GENE-6079153 | wingless-type MMTV integration site family, member 11B [Source:Xenbase;Acc:XB-GENE-5858981] | - | - | + |

| BMP pathway | | | | | |
|---|---|---|---|---|---|
| Genes | Xenbase.gene.ID | description | St 10 | St 11 | St 14 |
| bmp7.2.L | XB-GENE-856288 | bone morphogenetic protein 7, gene 2 [Source:Xenbase;Acc:XB-GENE-855954] | - | + | - |
| bambi.L | XB-GENE-866425 | BMP and activin membrane-bound inhibitor [Source:Xenbase;Acc:XB-GENE-495359] | - | - | + |
| bambi.S | XB-GENE-17343852 | BMP and activin membrane-bound inhibitor [Source:Xenbase;Acc:XB-GENE-495359] | - | - | + |
| bmp4.L | XB-GENE-483062 | bone morphogenetic protein 4 [Source:Xenbase;Acc:XB-GENE-483058] | - | - | + |
| bmp4.S | XB-GENE-6254046 | bone morphogenetic protein 4 [Source:Xenbase;Acc:XB-GENE-483058] | - | - | + |
| pnhd.L | XB-GENE-6488002 | pinhead [Source:Xenbase;Acc:XB-GENE-480588] | - | - | + |
| smad7.L | XB-GENE-865569 | SMAD family member 7 [Source:Xenbase;Acc:XB-GENE-486004] | - | - | + |

| Signaling (others) | | | | | |
|---|---|---|---|---|---|
| Genes | Xenbase.gene.ID | description | St 10 | St 11 | St 14 |
| ccn2.L | XB-GENE-855558 | cellular communication network factor 2 - growth factor [Source:Xenbase;Acc:XB-GENE-855547] | - | - | + |
| cxcr4.L | XB-GENE-489921 | C-X-C motif chemokine receptor 4 [Source:Xenbase;Acc:XB-GENE-489917] | - | - | + |
| igf3.L | XB-GENE-5807497 | insulin-like growth factor 3 [Source:Xenbase;Acc:XB-GENE-5807444] | - | - | + |
| mal.L | XB-GENE-5745230 | mal, T cell differentiation protein - Members of chemokine-like factor super family [Source:Xenbase;Acc:XB-GENE-5745206] | - | - | + |
| p2ry4.L | XB-GENE-943452 | pyrimidinergic receptor P2Y, G-protein coupled, 4 [Source:Xenbase;Acc:XB-GENE-943446] | - | - | + |
| sema3fl.S | XB-GENE-17335950 | semaphorin 3F like [Source:Xenbase;Acc:XB-GENE-940271] | - | - | + |
| tdgf1.2.S | XB-GENE-876243 | teratocarcinoma-derived growth factor 1, member 2 [Source:NCBI gene;Acc:100486054] | - | + | + |
| tab3.S | XB-GENE-6254217 | TGF-beta activated kinase 1 (MAP3K7) binding protein 3 [Source:Xenbase;Acc:XB-GENE-942652] | - | - | + |

| Extracellular matrix | | | | | |
|---|---|---|---|---|---|
| Genes | Xenbase.gene.ID | description | St 10 | St 11 | St 14 |
| chsy1.S | XB-GENE-17339959 | chondroitin sulfate synthase 1 [Source:Xenbase;Acc:XB-GENE-961883] | - | - | + |
| col5a2.L | XB-GENE-6485617 | collagen, type V, alpha 2 [Source:Xenbase;Acc:XB-GENE-6032476] | - | - | + |
| habp2.L | XB-GENE-6486520 | hyaluronan binding protein 2 [Source:Xenbase;Acc:XB-GENE-979150] | - | - | + |
| hapln3.S | XB-GENE-17346258 | hyaluronan and proteoglycan link protein 3 [Source:Xenbase;Acc:XB-GENE-5867355] | - | - | + |
| hyal2.L | XB-GENE-6464375 | hyaluronidase 2 [Source:NCBI gene;Acc:100144942] | - | - | + |
| mig30.L | XB-GENE-6251788 | Basement membrane-specific heparan sulfate proteoglycan (HSPG) core protein [Source:Xenbase;Acc:XB-GENE-6259003] | - | - | + |
| mmp3.L | XB-GENE-5932566 | matrix metallopeptidase 3 [Source:Xenbase;Acc:XB-GENE-5932509] | - | - | + |
| vcan.L | XB-GENE-922310 | versican [Source:Xenbase;Acc:XB-GENE-922302] | - | - | + |
| has-rs.S | XB-GENE-6252031 | hyaluronan synthase related sequence protein - Chitin synthase/hyaluronan synthase [Source:Xenbase;Acc:XB-GENE-12450674] | - | - | + |
| mmp28.L | XB-GENE-6254971 | matrix metallopeptidase 28 [Source:Xenbase;Acc:XB-GENE-984024] | - | - | + |
| spam1.L | XB-GENE-5793489 | sperm adhesion molecule 1 (PH-20 hyaluronidase, zona pellucida binding) [Source:Xenbase;Acc:XB-GENE-5793485] | - | - | + |
| timp2.S | XB-GENE-17334759 | TIMP metallopeptidase inhibitor 2 [Source:Xenbase;Acc:XB-GENE-949671] | - | - | + |

| Intercellular adhesion | | | | | |
|---|---|---|---|---|---|
| Genes | Xenbase.gene.ID | description | St 10 | St 11 | St 14 |
| cdh11.S | XB-GENE-864996 | cadherin 11 [Source:Xenbase;Acc:XB-GENE-483599] | - | - | + |
| cdh15.S | XB-GENE-6485759 | cadherin 15 [Source:Xenbase;Acc:XB-GENE-983662] | - | - | + |
| cldn4.L | XB-GENE-972446 | claudin 4 [Source:Xenbase;Acc:XB-GENE-972439] | - | - | + |
| cldn6.1.L | XB-GENE-959189 | claudin 6, gene 1 [Source:Xenbase;Acc:XB-GENE-959181] | - | - | + |
| dsp.S | XB-GENE-17346609 | desmoplakin [Source:Xenbase;Acc:XB-GENE-492195] | - | - | + |
| pcdh1.L | XB-GENE-983608 | protocadherin 1 [Source:Xenbase;Acc:XB-GENE-983603] | - | - | + |

| Cytoskeleton | | | | | |
|---|---|---|---|---|---|
| Genes | Xenbase.gene.ID | description | St 10 | St 11 | St 14 |
| arhgap42.L | XB-GENE-6487105 | Rho GTPase activating protein 42 [Source:NCBI gene;Acc:100493816] | - | - | + |
| arhgap45.L | XB-GENE-990083 | Rho GTPase activating protein 45 [Source:Xenbase;Acc:XB-GENE-990078] | - | - | + |
| cfap300.L | XB-GENE-943462 | cilia and flagella associated protein 300 [Source:Xenbase;Acc:XB-GENE-943457] | - | - | + |
| mfap2.L | XB-GENE-6254690 | microfibril associated protein 2 [Source:Xenbase;Acc:XB-GENE-6257614] | - | - | + |
| mtus1.L | XB-GENE-979441 | microtubule associated scaffold protein 1 XB | - | - | + |
| myo10.2.L | XB-GENE-17344263 | myosin 10, gene 2 [Source:Xenbase;Acc:XB-GENE-5995373] | - | - | + |
| mypn.L | XB-GENE-6486837 | myopalladin - Projectin/twitchin and related proteins [Source:Xenbase;Acc:XB-GENE-988987] | - | - | + |
| prph.L | XB-GENE-866295 | peripherin [Source:Xenbase;Acc:XB-GENE-493857] | + | + | + |
| rnd3.S | XB-GENE-17343479 | Rho family GTPase 3 [Source:Xenbase;Acc:XB-GENE-495412] | - | - | + |
| snx22.L | XB-GENE-17340049 | sorting nexin 22 [Source:Xenbase;Acc:XB-GENE-946153] | - | - | + |
| vim.L | XB-GENE-866225 | vimentin [Source:Xenbase;Acc:XB-GENE-493024] | - | - | + |
| cdc42ep2.L | XB-GENE-17345746 | CDC42 effector protein 2 [Source:Xenbase;Acc:XB-GENE-5818741] | - | - | + |
| ckap2.L | XB-GENE-6486301 | cytoskeleton associated protein 2 [Source:Xenbase;Acc:XB-GENE-1004436] | - | - | + |
| inf2.S | XB-GENE-5862221 | inverted formin, FH2 and WH2 domain containing [Source:Xenbase;Acc:XB-GENE-5862185] | - | - | + |
| myo5b.S | XB-GENE-6487867 | myosin VB [Source:Xenbase;Acc:XB-GENE-988439] | - | - | + |
| rhof.L | XB-GENE-17336827 | ras homolog family member F (in filopodia) [Source:Xenbase;Acc:XB-GENE-920536] | - | - | + |
| rhov.L | XB-GENE-17343382 | ras homolog family member V [Source:Xenbase;Acc:XB-GENE-489435] | - | - | + |
| rnd1.S | XB-GENE-17336490 | Rho family GTPase 1 [Source:Xenbase;Acc:XB-GENE-490045] | - | - | + |
| shroom2.L | XB-GENE-6485952 | shroom family member 2 [Source:Xenbase;Acc:XB-GENE-982291] | - | - | + |

| Channels/transporters | | | | | |
|---|---|---|---|---|---|
| Genes | Xenbase.gene.ID | description | St 10 | St 11 | St 14 |
| kcne1.L | XB-GENE-962321 | potassium channel, voltage gated subfamily E regulatory beta subunit 1 [Source:Xenbase;Acc:XB-GENE-962320] | - | - | + |
| slc12a1.L | XB-GENE-6486739 | solute carrier family 12 member 1 [Source:Xenbase;Acc:XB-GENE-479925] | - | - | + |
| slc12a3.L | XB-GENE-6488610 | solute carrier family 12 member 3 [Source:Xenbase;Acc:XB-GENE-866805] | - | - | + |
| slc1a5.S | XB-GENE-6077517 | solute carrier family 1 member 5 [Source:Xenbase;Acc:XB-GENE-482596] | - | - | + |
| slc2a2.L | XB-GENE-865613 | solute carrier family 2 member 2 XB | - | - | + |
| atp1b1.S | XB-GENE-864911 | ATPase Na+/K+ transporting subunit beta 1 [Source:Xenbase;Acc:XB-GENE-482813] | - | - | + |
| slc15a4.L | XB-GENE-986488 | solute carrier family 15 member 4 [Source:Xenbase;Acc:XB-GENE-986485] | - | - | + |
| slc38a2.L | XB-GENE-6254869 | solute carrier family 38 member 2 [Source:Xenbase;Acc:XB-GENE-6258676] | - | - | + |
| vdac1.S | XB-GENE-955070 | voltage-dependent anion channel 1 [Source:Xenbase;Acc:XB-GENE-955063] | - | - | + |

Proliferation/apoptosis

| Genes | Xenbase.gene.ID | description | St 10 | St 11 | St 14 |
|---|---|---|---|---|---|
| bok.S | XB-GENE-6251426 | BCL2 family apoptosis regulator BOK [Source:Xenbase;Acc:XB-GENE-988663] | - | - | + |
| dapk1.S | XB-GENE-6251779 | death associated protein kinase 1 [Source:Xenbase;Acc:XB-GENE-1011852] | - | - | + |
| cc-.L | XB-GENE-979551 | cyclin O [Source:Xenbase;Acc:XB-GENE-979545] | - | + | - |

Supplemental table 2: Genes differentially expressed upon RA disruption. Genes down-regulated that are expected to be positive RA targets are highlighted with green. Genes up-regulated that are expected to be negative targets are highlighted with red. Differential expression observed with a fdr < 0.05 and a $\log_2 fc > 0.5$ (absolute value) at a given stage is indicated by +. Manual annotation of some genes from Xenbase website pages information is indicated in red.

#### Supplementary Table 3. Scoring of meis3 depletion experiments.

| Gene expression analysis | Experiment | Momeis3 | | Control morpholino | |
|---|---|---|---|---|---|
| | | Affected left side | Normal left side | Affected left side | Normal left side |
| ISH *pax8* NFst18/19 | #1 | 9 | 8 | 4 | 17 |
| | #2 | 9 | 12 | 0 | 3 |
| | #3 | 18 | 5 | 2 | 13 |
| | #4 | 15 | 5 | 2 | 19 |
| | #5 | 16 | 4 | 3 | 19 |
| | **Total (%)** | **67 (66.3)** | **34 (33.7)** | **11 (13.4)** | **71 (86.6)** |
| ISH *lhx1* NFst18/19 | #1 | 9 | 7 | 4 | 15 |
| | #2 | 12 | 8 | 1 | 14 |
| | #3 | 19 | 2 | 2 | 14 |
| | #4 | 11 | 2 | 4 | 14 |
| | #5 | 18 | 2 | 5 | 16 |
| | **Total (%)** | **69 (76.7)** | **21 (23.3)** | **16 (18)** | **73 (82)** |
| ISH *aldh1a2* NFst18/19 | #1 | 6 | 5 | 2 | 17 |
| | #2 | 14 | 8 | 0 | 21 |
| | #3 | 13 | 2 | 4 | 10 |
| | #4 | 22 | 2 | 12 | 9 |
| | **Total (%)** | **55 (76.4)** | **17 (23.6)** | **18 (24)** | **57 (76)** |

A morpholino targeting meis3 (Momesi3) or a control morpholino were injected into the left V2 blastomere of 8-cell stage embryos. Expression of *pax8*, *lhx1* and *aldh1a2* was analyzed at late neurula stage (NFst18-19). Expression patterns corresponding to the representative examples shown in Figs 3 and 4 are scored for every independent experiment.

**Supplementary Table 4. Sequences of primers use for pax8-CNS1 mutations and RT-qPCR analyses.**

**Primers used for mutation of pax8-CNS1**

| **Name** | **Forward** | **Reverse** |
|---|---|---|
| pax8CNS1mut-lucMeis | 3’-CTCAGTAAACAGCCGCCGACTGTTGCATTAGCCTGGAAATGAATTCTCACTTTTATAGGG-5’ | 3’-CCCTATAAAAGTGAGAATTCATTTCCAGGCTAATGCAACAGTCGGCGGCTGTTTACTGAG-5’ |
| pax8CNS1mut-lucHP1 | 3’-GCCTGACACTGCATTAGCCTGGAAGCGGCGGGTCACTTTTATAGGGAAATTGCAAATC-5’ | 3’-GATTTGCAATTTCCCTATAAAAGTGACCCGCCGCTTCCAGGCTAATGCAGTGTCAGGC-5’ |
| pax8CNS1mut-lucHP2 | 3’-CACTGTCCCCGAGGGCGGATCCATGCTCAGTAAACAGCCGCCTG-5’ | 3’-CAGGCGGCTGTTTACTGAGCATGGATCCGCCCTCGGGGACAGTG-5’ |
| pax8CNS1mut-lucHP3 | 3’-GAGGAGTCAGGATTTAATTCAGCGGGCGGTTAACTAGTCCTCTGTATGCACTGTCCCC-5’ | 3’-GGGGACAGTGCATACAGAGGACTAGTTAACCGCCCGCTGAATTAAATCCTGACTCCTC-5’ |

**Primers used for RT-qPCR**

| **gene** | **Forward** | **Reverse** |
|---|---|---|
| *ak1b* | 3’-CAGCCATCATGGAGAGAGGAG-5’ | 3’-TGTCCAGAACTGTGTCCAAAGG-5’ |
| *aldh2a1* | 5'-TTCAGCAATGCAAGCAGGAA-3' | 5'-GCTTTGAGCATTCAGAGCATTGTA-3' |
| *beta actin* | 3’-TCTATTGTGGGTCGCCCAAG-5’ | 3’-TTGTCCCATTCCAACCATGAC-5’ |
| *cyp26a1* | 5'-GGAGACTCTGCAAATGGTGCT-3' | 5'-CGCTTCATTTGGAGGAACTTG-3' |
| *hoxa1* | 5'-CCACCCAAAACAGGTAAGGCT -3' | 5'-GTTGGGCTGGCCAACATAA-3' |
| *hoxa3* | 3’-ACAAGAGGTCGGACCAGGG-5’ | 3’-TCTGTCACGTGGCGTTCCT-5’ |
| *hoxb3* | 3’-AAGAAGCCCCCAAATTGACC-5’ | 3’-GAGGTGATGGTTGGGCTGAG-5’ |
| *hoxb4* | 3’-CCGTGGTGTACCCCTGGAT-5’ | 3’-TGGAAGTGGCCTTGGAGATG-5’ |
| *hoxc5* | 3’-ACCAAACTGCACATGAGCCA-5’ | 3’-ACTTGTCCGGGAACGCTTC-5’ |
| *Hoxd1* | 3’-GTCAGCACGTTTGATTGGATGA-5’ | 3’-TTTGTCCTCACATTGCAGGG-5’ |
| *lhx1* | 5'-CAACATGCGAGTAATCCAGGTC-3' | 5'-CTCTCTTTGGATCGTCGGTTCT-3' |
| *meis3* | 3’-TACGATTCCACCTGCTGGAGT-5’ | 3’-CGGTGGCAGAAATTGTCACAC-5’ |
| *pax8* | 5’- CAGCAATTTCAATATAGGTCACGG-3’ | 5’- TCCATTCACAAAAGCCCCAC-3’ |
| *odc* | 3’-GGGCAAAGGAGCTTAATGTGG-5’ | 3’-TGCCAACATGGAAACTCACAC-5’ |
| *sema3f* | 3’-AAGCAGGAAAACATGGCGAC-5’ | 3’-CACGCCTGCATACAGCACTC-5’ |
| *sh3bp5* | 3’-GATTTGATTGGTCTGCCCTTG-5’ | 3’-CCCAGATCAGCAACCTTCCTC-5’ |
| *slc12a1* | 3’-ACCGCCAAGTCAGACTGAATG-5’ | 3’-TTGCCACAGGAAGACTCAGGA-5’ |
| *tmem38a* | 3’-CATGGTACTGATCGGATGGGT-5’ | 3’-TTGGACATCAGGGCCACTC-5’ |
| *vcan* | 3’-GGGTACAAGGCGGTCACCT-5’ | 3’-CTTATCGTTGAGGCCAATCCA-5’ |

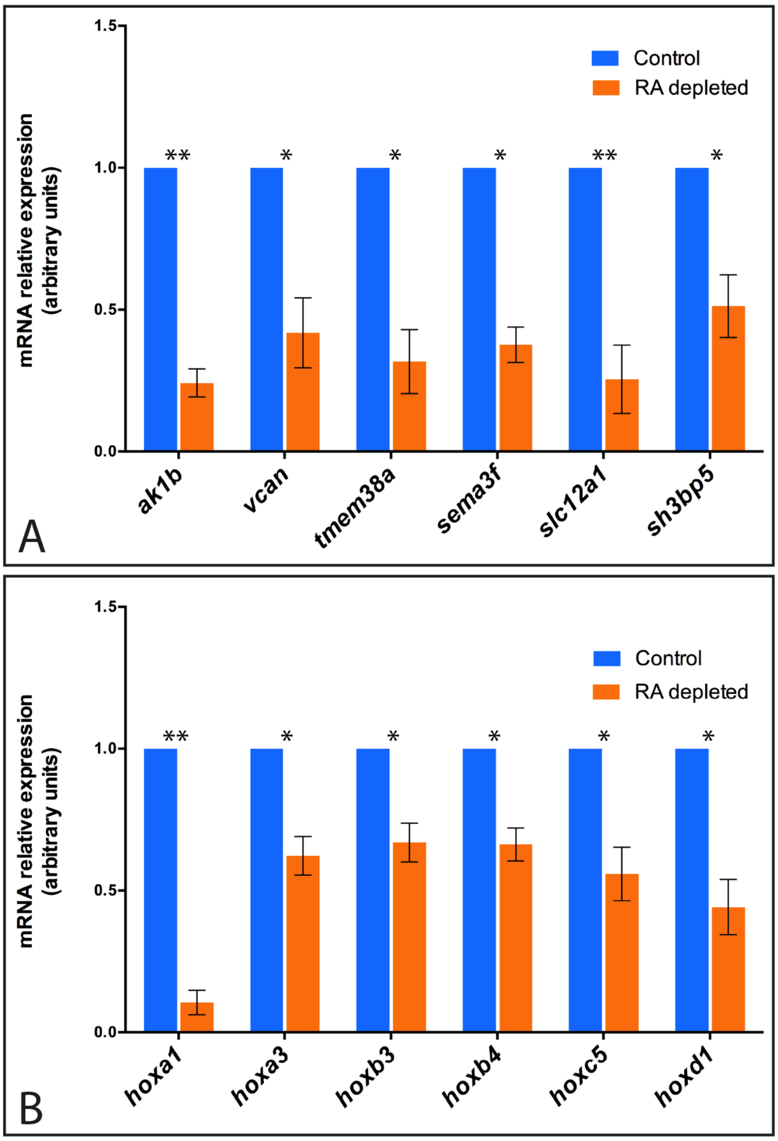

Supplemental Figure 1

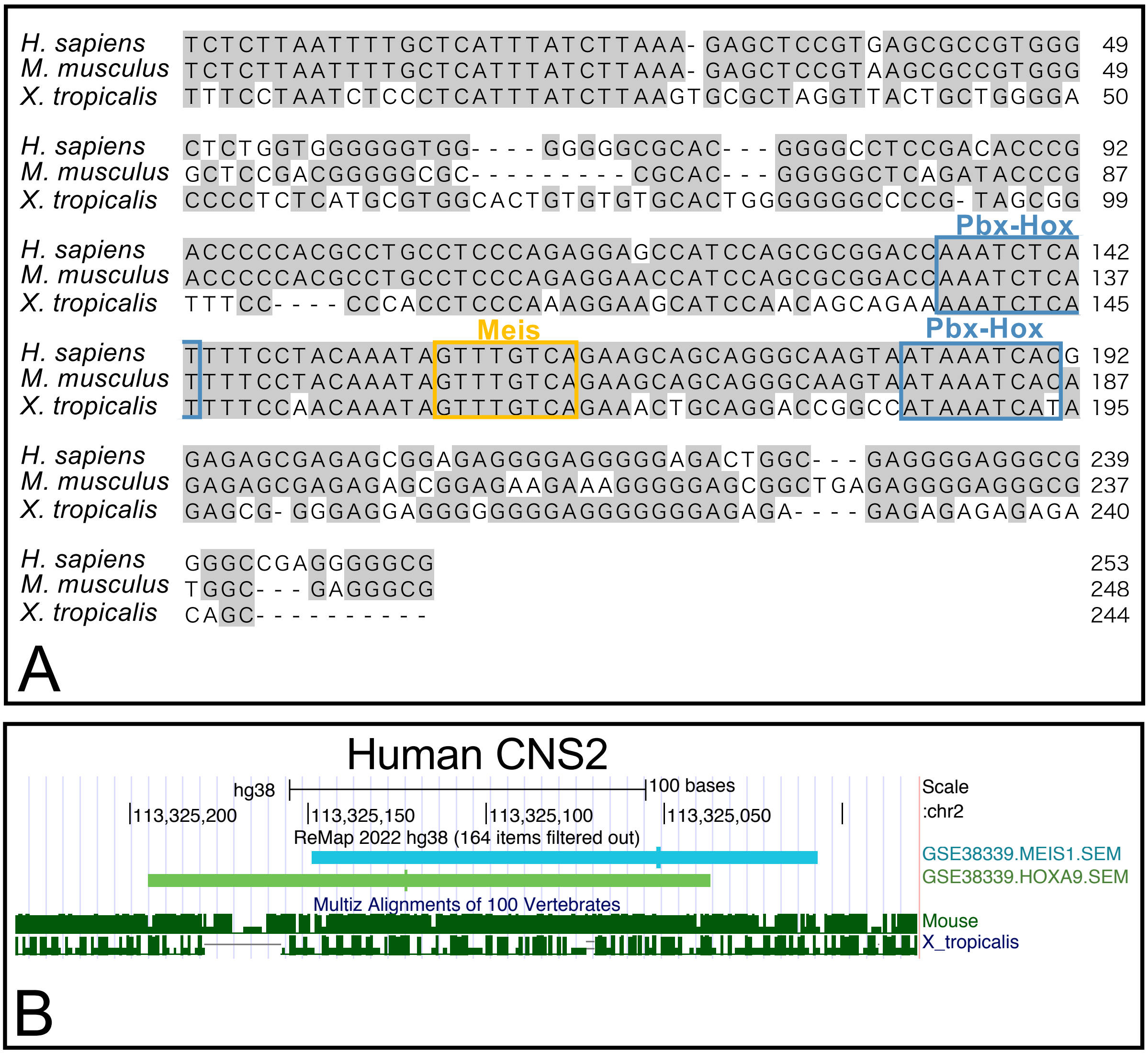

Supplemental Figure 2
